# Negative allosteric modulation of α5-GABA_A_ receptors engages dynamic cortical glutamatergic and GABAergic mechanisms underlying adaptive behavior in mice

**DOI:** 10.64898/2026.01.15.699507

**Authors:** Fernanda Daher, Caio T. Fukushima, Erik A. Ingebretsen, Jean M. Bidlack, John R. Atack, Mariana O. Popa, Manoela V. Fogaça

## Abstract

Chronic stress disrupts glutamatergic and GABAergic plasticity in the medial prefrontal cortex (mPFC), impairing circuit integration and contributing to the pathophysiology of stress-related disorders, such as Major Depressive Disorder (MDD). Rapid-acting antidepressants like ketamine can rapidly reverse these deficits, but its clinical use is limited by psychotomimetic side effects. Notably, the α5-GABA_A_R negative allosteric modulator (α5-NAM) Basmisanil (BSM), reproduces ketamine-like behavioral outcomes in preclinical models, although the cellular mechanisms underlying its actions remain unclear. Here, we investigated whether BSM promotes ketamine-like enhancement of cortical plasticity and engages cell type-specific mechanisms to support adaptive behaviors over time. We show that BSM produced rapid and sustained facilitation of motivational, hedonic, and active coping behaviors via mPFC circuits. BSM induced c-Fos expression in mPFC D1R- and somatostatin-expressing cells, suggesting activation of specific subsets of pyramidal and GABA interneurons. In both mPFC and hippocampus, BSM rapidly activated Erk- or Akt-mTOR signaling pathways as well as increased synaptic proteins critical for glutamatergic and GABAergic function. BSM also reversed maladaptive behaviors induced by chronic unpredictable stress, including impairment in object recognition memory and social interaction. Finally, chemogenetic silencing of mPFC CaMKII-expressing neurons blocked both rapid and sustained actions of BSM, whereas inhibition of mPFC GABA interneurons reversed only long-term behavioral outcomes. These results indicate that α5-GABA_A_R modulation requires early activation of pyramidal neurons to drive rapid plasticity, while GABAergic adaptations support sustained improvements. This dynamic mechanism restores excitation–inhibition (E/I) balance and highlights GABAergic pathways as therapeutic targets for prefrontal dysfunction in stress disorders.

## INTRODUCTION

Among prevalent mental health conditions, Major Depressive Disorder (MDD) accounted for approximately 332 million cases worldwide in 2021 (1), and represented 37.3% (the largest proportion) of disability-adjusted life-years in 2019 (2). In addition to its significant global burden, the persistence of treatment-resistant or difficult-to-treat depression ratifies the urge for new evidence-based interventions (3, 4). Yet, progress in psychiatric drug development remains slow: in 2024, only one out of 50 new molecular entities approved by the FDA’s Center for Drug Evaluation and Research was a psychiatric drug (5), highlighting that despite remarkable advances in drug discovery, new psychiatric treatments have not progressed as in other therapeutic areas.

Ketamine is the prototypic rapid-acting antidepressant in clinical and preclinical studies. While intravenous racemic ketamine remains an off-label treatment, the intranasal *S*-enantiomer of ketamine (esketamine) has been FDA approved for treatment-resistant depression (6). In humans, ketamine increases glutamate and GABA metabolites in the prefrontal cortex (PFC) and restores PFC impaired connectivity in MDD patients during infusion and up to 24 h afterward (7–9). Preclinical studies suggest that ketamine acts by disinhibiting glutamate release through blockade of NMDA receptors on GABAergic interneurons and/or by directly blocking these receptors postsynaptically (10–12). Both indirect and direct mechanisms converge to stimulate AMPA glutamate receptors, raise BDNF levels and enhance synaptic efficacy in the medial PFC (mPFC). Following these fast mechanisms, ketamine promotes adaptations in GABAergic interneurons supporting sustained behavioral recovery (9, 13, 14). However, ketamine administration is associated with transient psychotomimetic effects (15, 16), which contribute to its potential for misuse and addiction (17), representing a major clinical concern.

Growing evidence highlights the GABAergic system as a promising therapeutic target in depression, particularly since GABAergic deficits are highly implicated in MDD’s pathophysiology (18, 19). Within the functional diversity of type A γ-aminobutyric acid receptors (GABA_A_Rs), α5 subunit-containing GABA_A_Rs (α5-GABA_A_Rs) exhibit distinctive functional properties. First, α5-GABA_A_Rs are densely enriched in the PFC and hippocampus of rodents and humans (20–22), brain regions critical for emotional regulation, sociability, decision-making and memory formation. Second, α5-GABA_A_Rs are predominantly expressed extrasynaptically, where it mediates tonic, persistent inhibition in response to ambient GABA (23, 24). Further, incorporation of the α5 subunit into the pentameric ligand-gated ion channel in diverse arrangements generates specific allosteric binding sites sensitive to pharmacological modulation (25, 26). Paradoxically, both an increase or a decrease of the apparent affinity of extrasynaptic GABA_A_R for GABA by positive or negative allosteric modulators (PAMs or NAMs, respectively) is shown to result in antidepressant effects (27–29). Notably, the neuroactive steroids brexanolone and zuranolone, which act as PAMs of γ-GABA_A_R, have been approved for the treatment of postpartum depression in adult women (30) since 2019 and 2023, respectively, offering a mechanistically distinct alternative to traditional antidepressants developed decades ago that target serotonin, dopamine and/or norepinephrine systems.

On the other side of this bidirectional pharmacological spectrum, α5-GABA_A_R NAMs (α5-NAMs) have been primarily linked to memory enhancement and, more recently, to antidepressant-related effects in preclinical assays (27, 28, 31). MRK-016, an α5-NAM, enhances cognitive performance in rodents (32), reverses stress-induced anhedonia and deficits in social interaction in mouse models of chronic stress (28, 33), and restores excitatory synaptic strength in hippocampal slices (28, 31, 33). However, clinical development of MRK-016 was halted due to variable interindividual pharmacokinetics, and poor tolerability issues (34). In contrast, Basmisanil (BSM), another α5-NAM, has demonstrated a safety profile in phase II clinical trials evaluating its efficacy as a cognitive enhancer (35, 36). BSM combines subtype-selective affinity (100−200-fold for α5- over α1-, α2-, and α3-subunits) and α5-NAM efficacy (34, 36), which may minimize off-target effects while reducing GABAergic dendritic inhibition of pyramidal cells in the PFC.

Notably, our recent findings report rapid and sustained antidepressant-like behavioral effects in male mice under low-to-moderate levels of estimated α5 occupancy following 10-mg/kg BSM administration (27), suggesting that BSM may reverse structural and behavioral deficits within stress-sensitive circuits. However, it is unknown whether a single injection of BSM exerts fast and sustained antidepressant-like effects via convergence on ketamine’s downstream signaling pathways, and how glutamatergic excitatory and GABAergic inhibitory neurons contribute to these actions. Here, we investigate the behavioral, synaptic, and cellular changes induced by BSM using well-established pharmacological, molecular, and chemogenetic approaches in male mice. Our converging evidence supports the hypothesis that α5-NAMs rapidly enhance glutamatergic drive and activity-dependent intracellular plasticity mechanisms in the mPFC, with long-lasting GABAergic adaptations supporting behavioral recovery.

## MATERIALS AND METHODS

### Animals

Male C57BL/6 mice (8-14-week-old) were obtained from The Jackson Laboratory (B6J, strain No. 000664) or in-house breeders. Calcium-calmodulin-dependent protein kinase II alpha (*CaMKIIa)*^Cre^ or glutamic acid decarboxylase 2 (*Gad2*)^Cre^ transgenic male mice, and respective WT littermates (8-14-week-old) on the C57BL/6 background (donated by the Picciotto’s lab, Yale University, originally obtained from Jackson) were bred in-house as reported (37). All animals were group-housed in ventilated home cages (29 cm length × 18 cm width × 13 cm height) with a 12/12h light-dark cycle, food, and water *ad libitum*, except when social isolation, inverted cycle and/or water deprivation was used as stressors. Male mice were used in this study to facilitate direct comparison with the existing ketamine literature, which has been conducted predominantly in males; future studies will include female mice. All procedures were approved by the Institutional Animal Care and Use Committees (IACUC) of the University of Rochester and conducted per the guidelines for the care and use of laboratory animals of the National Institutes of Health (NIH).

### Surgical procedures

*Cannula placement*: C57BL/6 mice were anesthetized (ketamine 100 mg/kg and xylazine 10 mg/kg, intraperitoneally, i.p.) and received implantation of bilateral guide cannulas (part No. C235GS; 26G, 1-mm center-to-center distance and 7-mm total length; Protech International Inc., Boerne, TX, USA). Cannulas were implanted in the mPFC 0.2 mm above the microinjection site (stereotaxic coordinates relative to bregma: +1.9 mm anterior; ±0.4 mm lateral; -1.7 mm ventral). Dental cement (DuraLay Standard Package, Reliance Dental Manufacturing, Alsip, IL, USA) maintained the guide cannulas in place, and 2 screws (2-mm length and 1-mm diameter) inserted into the skull anchored the dental cement. After 8-day recovery, mice received bilateral microinjection (0.2 µL/side; 0.2 µL/min) into the mPFC using injectors (33G) extending 1 mm below the guide cannulas.

*AAV infusion*: Adeno-associated viruses AAV2-hSyn-DIO-hM4D(Gi)-mCherry (hM4D(Gi), ≥ 5 x 10^12^ vg/ml), and AAV2-hSyn-DIO-mCherry (mCherry, ≥ 4×10¹² vg/mL) were obtained from Addgene (USA). Anesthetized CaMKII^Cre+^ or GAD1^Cre+^ mice and WT littermates received bilateral microinjection of inhibitory hM4D(Gi) into the mPFC (0.5 μL/side; 0.1 μL/min; aforementioned coordinates). For analgesia, animals received subcutaneous injections of carprofen (5 mg/kg) immediately before the surgery, 24 and 48 h post-operation.

### Drug administration

*Drug injection (i.p.)*: BSM (10 mg/kg i.p.; Medicines Discovery Institute at Cardiff University; PubChem CID 57336276) was microsuspended in Tween® 80 2% (v/v) in 0.9% sodium chloride (saline). The vehicle (control) group received Tween® 80 2% in saline at the same dose volume. Ketamine (10 or 20 mg/kg, i.p.; Sigma-Aldrich) was dissolved in saline. Behavioral testing started 1 h after treatment and continued for up to 48-50 h; perfusion for c-Fos immunofluorescence was performed 1.5 h after treatment, and euthanasia for Western Blotting was conducted 1 or 24 h post-treatment.

*Intra-brain drug microinjection*: BSM (3; 10; or 30 ng, intra-mPFC, 0.2 µL/side) was microsuspended in dimethyl sulfoxide (DMSO) 10% (v/v) in saline. The vehicle group received DMSO 10% in saline. Behavioral testing started 30 min after treatment and continued for up to 48 h.

*Corticosterone (CORT) administration:* CORT (Sigma-Aldrich) was dissolved in 1% ethanol (EtOH) in Milli-Q water and administered via the drinking water at a concentration of 0.1 mg/mL for 21 days (38). Control mice received water only.

*Chemogenetic inhibition*: Clozapine-N-oxide (CNO, 1 mg/kg i.p.; Enzo Life Sciences, Farmingdale, NY, USA) was dissolved in saline. Our previous studies indicate that CNO at 1 mg/kg does not produce behavioral or locomotor effects *per se* (39). All CaMKII^Cre+^, Gad^Cre+^, and WT groups received CNO in this study to control for potential off-target effects associated with clozapine conversion at higher doses (5–10 mg/kg) (40, 41). CNO was administered 20 min before BSM or vehicle, equivalent to 80 min before the onset of behavioral testing.

### Chronic Unpredictable Stress (CUS)

CUS is a chronic stress paradigm to induce maladaptive behaviors that recapitulates reminiscent of psychiatric symptoms (42), which are reversed by a single injection of ketamine (43). In the present study, mice were exposed to a sequence of two random unpredictable stressors per day for 21 or 28 days, as we have described (39, 44, 45). Cohorts tested in the behavioral battery underwent 21-day CUS; cohorts tested in the social interaction (SI) test underwent 28-day CUS, in which mice remained isolated for 21 out of 28 days to promote social deficits. A variety of 12 stressors was applied to avoid habituation (Table 1) (14, 39, 44), and exposure to stressors occurred at least 3 h apart. To avoid acute effects of stress, mice were not exposed to any stressors on the day of treatment or behavioral testing. Stressed mice were divided into three groups and received a single i.p. injection of vehicle, ketamine or BSM. Non-stressed mice were handled weekly but not exposed to stressors and received a single i.p. injection of vehicle.

**Table 1.**
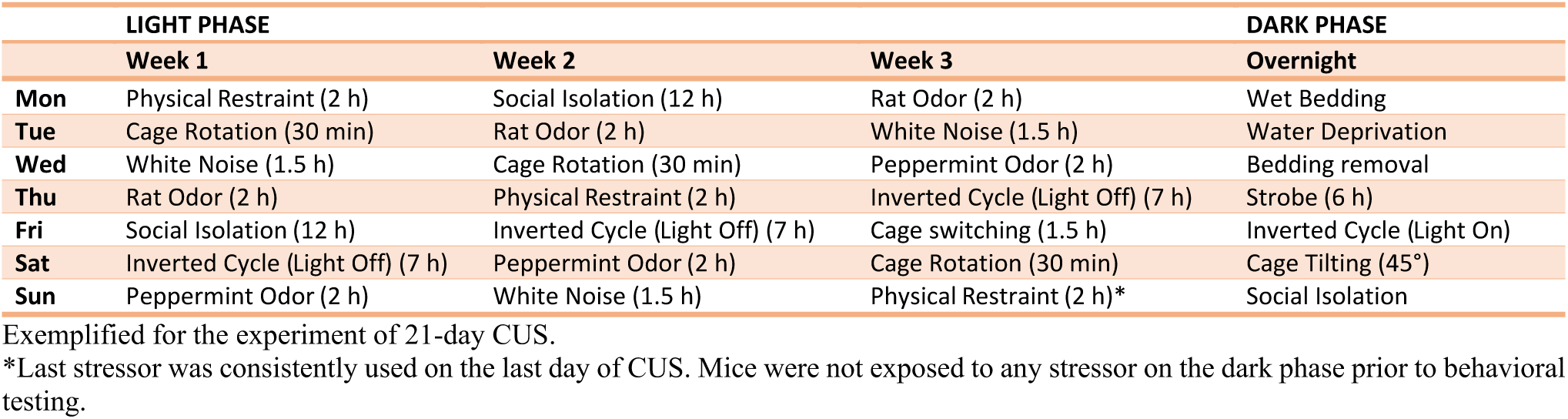
Chronic unpredictable stress (CUS) schedule.

### Behavioral studies

Before each behavioral assay, mice were left undisturbed in the testing room at least 30 min for acclimatization. Tests were performed between 9 a.m. and 3 p.m. Analyses were performed by an experimenter blind to treatment conditions whenever possible.

*Forced Swim Test (FST)*: Adult mice were placed in a clear 5-L beaker filled with water (26°C ± 1; 300 lux ambient lighting) and sessions were scored for total immobility time during minutes 2 to 6, as previously described (39, 44). Mice underwent two swimming sessions: a pre-swim session (before treatment) to establish a stable immobility baseline for the following day, and a test session 24 h later (after treatment). In experiments involving CUS, mice underwent only the test session to avoid a ceiling effect.

*Open Field Test (OFT)*: To assess locomotor activity during exploration of a novel environment, mice were placed into individual open-field arenas (40 cm length × 40 cm width × 30 cm height; Maze Engineers, Skokie, IL, USA; 150 lux ambient lighting) for 10 min (46). Distance traveled, and speed were analyzed using EthoVision XT 17.5.1718 (Noldus Information Technology Inc., Leesburg, VA, USA).

*Sucrose Splash Test (SuST)*: To examine self-care motivation and response to a natural reward, mice were singly placed in standard home cages (29 cm length × 18 cm width × 13 cm height; red ambient lighting) without bedding and a fresh 10% sucrose solution was squirted onto the dorsal coat, as described (39, 44, 47). Grooming time was measured for 5 min.

*Female Urine Sniffing Test (FUST)*: To assess natural reward–seeking and hedonic behavior, each mouse was placed in a freshly prepared individual home cage for a 50-min habituation period to a sterile cotton-tipped applicator (10 lux ambient lighting) (44). The applicator was inserted through a small hole in the cage lid and secured with a clip. After habituation, the applicator was removed, and the mouse was first presented with a new cotton-tipped applicator dipped in tap water (control) for 5 min. This was then replaced with a fresh applicator containing female urine for an additional 5 min. Sniffing time toward the water and urine stimuli was quantified over their respective 5-min periods; time spent biting or climbing the applicator was not scored.

*Tail Suspension Test (TST)*: Mice were suspended by the tail using adhesive tape affixed to a shelf under 300 lux ambient lighting. Clear hollow cylinders were placed around the tails of the mice to prevent tail climbing behavior (48). Immobility time was measured over 6 min.

*Novel Object Recognition (NOR) Test and Re-Test*: To assess recognition memory that can be impaired by chronic stress and reversed using pharmacological treatments in adult rodents (49, 50), the NOR task was carried out over 4 consecutive days. Mice were first habituated to an empty open-field arena (20 lux ambient lighting) and, 24 h later, to the same arena with 2 identical objects (50-mL tissue culture flasks filled with white sand). On day 3, mice explored the familiar objects and, after a 3-h delay, were tested with one familiar and one novel object (Lego bricks). At day 4, mice were re-tested with a distinct novel object (either a Rubik’s cube or plastic lamp filled with white sand). Object exploration was quantified as time spent sniffing each object over a 10-min period, and recognition memory was indexed by a discrimination ratio for novel object preference: *(time exploring novel − time exploring familiar)/(time exploring novel + time exploring familiar)*.

*Social Interaction (SI) Test*: To assess interest in social novelty (51), the SI test was performed over 3 subsequent days. Experimental mice were habituated to an open-field arena for 10 min under 20 lux illumination. After 24 h, mice were placed in the same arena with an empty cylindrical barred cage positioned along one wall for 10 min. On day 3, mice underwent a second 10-min familiarization session (empty target) followed by the test session in which an unfamiliar adult male mouse (social target) of the same sex, strain and age was placed inside the barred cage for 10 min (with a 6-min inter-session interval to clean the arena). Total investigation time was scored during the second familiarization session and the test session to quantify empty-enclosure exploration and sociability, respectively.

*Drug-paired conditioned place preference (CPP):* Mice were placed in a CPP apparatus, as previously described (52, 53). The apparatus consisted of two outer compartments (27.3 × 22.2 × 34.9 cm), each with distinct visual (vertical vs. horizontal black-and-white stripes, 1.5 cm width) and tactile (lightly mottled vs. smooth floor) cues, connected by a smaller central chamber (13.9 × 22.2 × 34.9 cm) with sliding doors (Place Preference, San Diego Instruments, San Diego, CA). Infrared beams along the walls allowed automated measurement of time spent in each compartment and ambulatory activity. Mice were habituated to the apparatus for two consecutive days (20 min/day) with all doors open. On day 2, the outer compartment in which a mouse spent the most time was designated as its initially preferred context (baseline CPP or pre-CPP). Conditioning began the following day: mice received drug or vehicle and were confined to their initially non-preferred chamber for 30 min. Twenty-four hours later, all groups received saline and were placed in their initially preferred chamber for 30 min. This pairing sequence (drug or vehicle followed 24 h later by saline) was repeated three times over six days (days 3 to 8). Place preference was assessed on day 9 (54). CPP differences were computed as Δ time (*paired − unpaired*) within each session (pre-CPP and post-CPP). For graphical representation that allows visual comparison with the sucrose-paired CPP, a constant was added to all values to yield only positive numbers, and data were normalized to the vehicle pre-test condition (%). Statistical analyses were performed on untransformed data. Locomotor activity was assessed on the first conditioning day for 20 min immediately following drug or vehicle administration.

*Sucrose-paired CPP:* We adapted and implemented a three-arm T-maze paradigm to assess sucrose-paired conditioned place preference (CPP). Unlike conventional two-compartment sucrose CPP assays (55), this design incorporates a third, unpaired start arm of equal dimensions, enabling discrimination among reward-associated, non-rewarded, and neutral contexts. This added task complexity provides a more stringent assessment of goal-directed choice rather than simple binary preference and enhances sensitivity to motivational salience and executive control processes relevant to prefrontal cortical function. Group-housed mice received CORT in the drinking water for 21 days, while control mice received water only. Animals were weighed daily throughout the 10-day behavioral protocol. Prior to behavioral testing, mice were transitioned to single housing with normal drinking water and *ad libitum* food. To prime sucrose consumption, eight sucrose pellets were placed in the home cage for four days. Mice were then food-restricted to 3 g chow per day for the remainder of the experiment to enhance reward salience. On Day 1 (habituation), mice freely explored the T-maze for 15 min. On Day 2 (baseline or pre-CPP), mice were given full access to all arms for 15 min with video tracking to establish initial arm preference. During Days 3–6 (conditioning), mice underwent two daily sessions (morning and afternoon), during which access was restricted to two arms per session (the start arm and either the paired or unpaired arm). Sucrose (8 pellets) was consistently paired with the initially non-preferred arm, whereas the preferred arm was never paired with reward. Both the paired and unpaired arms contained distinct spatial contextual cues, while the third arm, which consistently served as the start arm, remained neutral and unpaired throughout conditioning. Because the start arm was never associated with reward or contextual cues, its novelty tend to habituate and decrease across sessions, whereas the unpaired arm, despite lacking reward, retains contextual salience through repeated exposure, enabling discrimination among neutral, non-rewarded, and reward-associated contexts. On Day 7, mice received an intraperitoneal injection of ketamine, BSM, or saline, 24 h prior to testing. On Day 8, CPP expression was assessed during a 15-min free-access session. For sucrose CPP, differences were calculated as described above; however, the unpaired condition was defined as the combined time spent in the start arm and the unpaired arm.

### Viral efficiency and histology check

Following the end of behavioral testing, mice were euthanized to check the efficiency of cannula placement or viral infusion. Mice were deeply anesthetized with an overdose of xylazine + ketamine solution (i.p.), and perfused with phosphate-buffered saline (PBS) and 4% paraformaldehyde (PFA). Brains were harvested, post-fixed in 4% PFA followed by cryoprotection in 30% sucrose at 4 °C for at least 48 h. Coronal sections were cut on a cryostat (30 or 60 μm). Microinjection sites were checked (Fig. 1G) using an ECHO Revolve microscope (model RVL-100-G, Discover Echo Inc., San Diego, CA, USA). Cre-dependent expression of hM4Di-mCherry in mPFC- containing slices were checked using a fluorescence microscope (Keyence BZ-X800; IL, USA).

**Figure 1.**
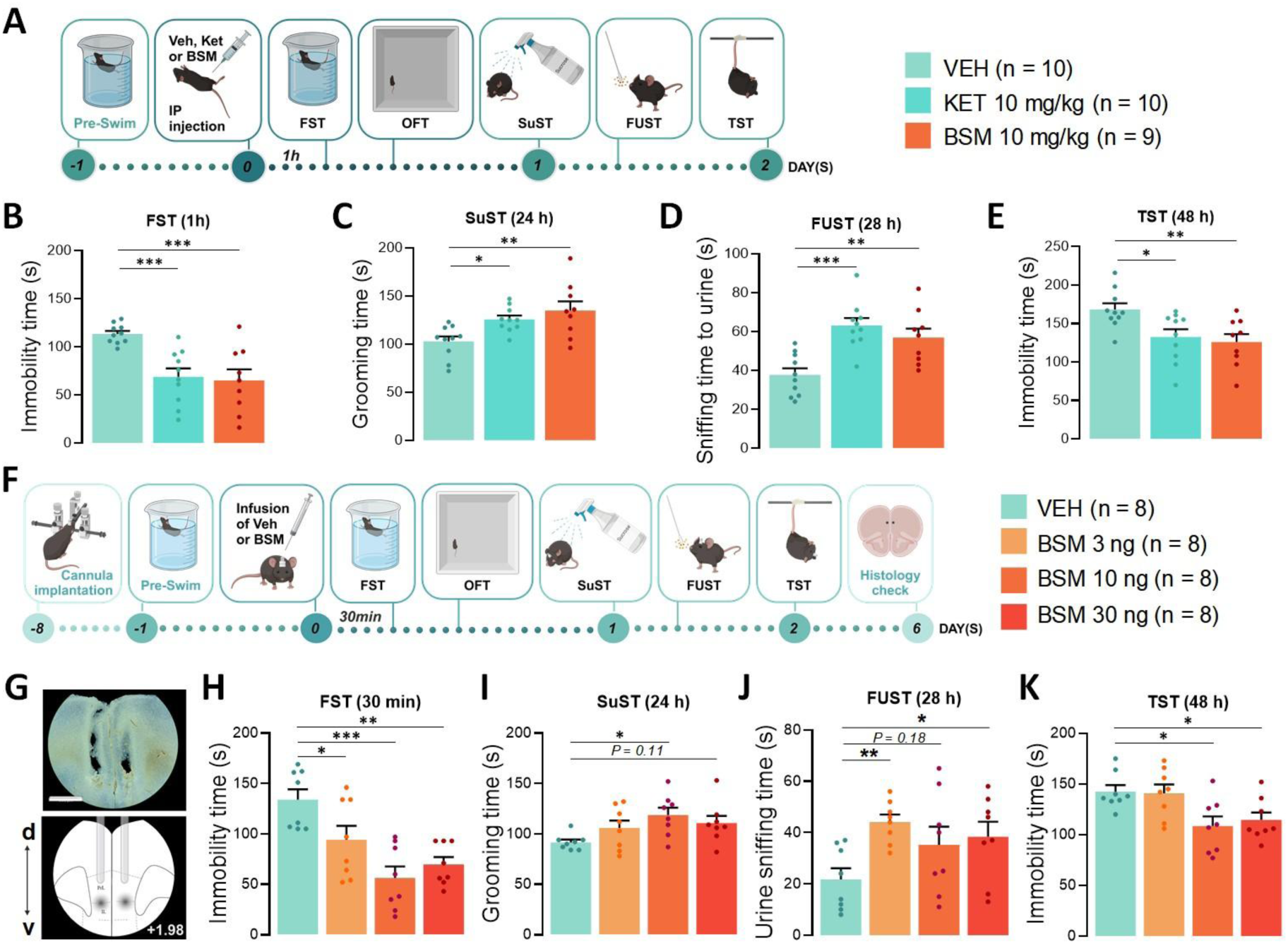
Basmisanil (BSM) facilitates rapid and sustained motivational, hedonic and active coping behaviors through mPFC modulation in male mice. (**A**) Schematic of behavioral test battery performed 1, 24, 28 and 48 h after a single injection of vehicle (VEH, negative control), ketamine (KET 10 mg/kg, i.p., positive control) or BSM (10 mg/kg, i.p.). (**B**) Average time spent immobile during the forced swimming test (FST, H(2) = 14.51; P < 0.001), (**C**) self-grooming during the sucrose splash test (SuST, F(2, 26) = 6.43, P = 0.005), (**D**) sniffing a cotton tip dipped in female urine during the female urine sniffing test (FUST, F(2, 26) = 11.56; P < 0.001), or (**E**) immobile during the tail suspension test (TST, F(2, 26) = 5.75; P = 0.009). (**F**) Mice were implanted with a bilateral guide-cannula above the mPFC and microinjected with VEH or BSM (3, 10, or 30 ng; 0.2 μL/side). The behavioral test battery was performed 30 min and 24, 28 and 48 h post-microinjection. (G) Image showing the cannula location and microinjection sites at 4× magnification in a mouse (top) and a schematic at bregma +1.98 (bottom; scale bar: 890 μm). (H) Behavioral changes were measured in the FST (H(3) = 16.83; P < 0.001), (I) SuST (F(3, 28) = 3.10; P = 0.04), (J) FUST (F(3, 28) = 3.26; P = 0.036) and (K) TST (F(3, 28) = 4.79; P = 0.008). Data are plotted as mean ± SEM. P values were calculated using independent-samples Kruskal-Wallis test or one-way ANOVA with Dunnett’s multiple-comparisons test. *P ≤ 0.05; **P ≤ 0.01; ***P ≤ 0.001. Created with BioRender.

### Immunofluorescence

Mice were deeply anesthetized with isoflurane (inhalation) and then transcardially perfused as aforementioned; brains were subsequently sectioned coronally at 30 µm thickness. Primary antibodies against c-Fos (rabbit anti-c-Fos #ab214672, 1:1000, Abcam), VGAT (rabbit anti-VGAT #131002, 1:1000, Synaptic Systems), VGLUT1 (guinea pig anti-VGLUT1 #135304, 1:2000. Synaptic Systems), dopamine type 1 receptor (D1R, rat anti-D1R #D2944, 1:500, Sigma-Aldrich), somatostatin (rat anti-SST #MAB354, 1:100, EMD Millipore), and parvalbumin (mouse anti-PV #AB277625, 1:1000, Abcam) were used (14, 39, 44). Appropriate secondary antibodies (1:1000, Millipore) were conjugated with Alexa Fluor® 488, 594 or 647. Sections were mounted onto slides and coverslipped. For c-Fos analysis, images containing the mPFC (prelimbic, PL or infralimbic, IL) or hippocampus (ventral, vHC or dorsal, dHC) were acquired using a fluorescence microscope at 4× or 10× magnification (Keyence BZ-X800). Regions of interest were defined manually based on anatomical landmarks. Total number of c-Fos^+^ cells in compressed z-stack images were quantified using ImageJ software and normalized by area, as we have described (39, 44). Number of c-Fos^+^ cells/area were obtained within each region of interest and averaged per mouse (3-4 slices/mouse). The results were then averaged across groups (n = 4-5 mice). In each experiment, the same exposure conditions (laser, gain and offset) were used for all slices. To determine the percentage of c-Fos^+^ cells that express D1R, SST or PV, a colocalization analysis was performed. For this, z-stack images containing the IL area were taken with a 40×-objective, and co-localized cells were manually counted by an experimenter blind to treatment conditions. Results are expressed as (*number of co-localized cells)/(number of total c-Fos) × 100*. For VGLUT1 and VGAT expression analysis, compressed z-stack images (20×) containing the PL or IL were analyzed with ImageJ to measure the integrated density for each photo. Values obtained from 3–4 sections were averaged for each animal, and results were expressed as group means normalized to the vehicle-treated group (%), as we have described (39, 44).

### Western blotting

Western blotting was conducted as previously described (39, 44, 56, 57), using the synaptoneurosome fraction of mPFC or hippocampal homogenates. Briefly, tissues were collected into RIPA lysis buffer (50mM Tris–HCl pH 7.5, 150mM NaCl, 1% Triton X-100, 0.1% SDS, 1mM NaVO3, 5mM NaF and 1× protease inhibitor cocktail). Tissues were then homogenized in a solution containing 0.32M sucrose, 20mm Hepes (pH 7.4), 1mM EDTA, 1× protease inhibitor cocktail, 10mM NaF and 1mM NaVO3. Homogenates were centrifuged at 2,800 rpm, supernatants were collected and further centrifuged at 12,000 rpm to obtain a pellet containing crude synaptoneurosomes. Pellets were then resuspended and sonicated in RIPA lysis buffer. Total protein concentrations were measured using a Pierce BCA Assay kit (ThermoFisher Scientific, USA). Proteins were electrophoretically separated on a precast gel (Criterion TGX, Bio-Rad Laboratories, Inc., USA) and transferred to a PVDF membrane. Proteins were detected with primary antibodies previously validated (39, 44, 57) and relevant to: (1) mammalian target of rapamycin (mTOR) signaling pathway: rabbit anti-phospho-Ser/Thr kinase AKT, #4058S, 1:1000; anti-AKT #9272S, 1:1000; anti-phospho extracellular signal-regulated kinase (ERK 1/2), #9101S, 1:1000; anti-ERK 1/2 #9102S, 1:2000; anti-phospho-glycogen synthase kinase-3β (pGSK-3β), #9336S, 1:500; anti-GSK-3β #9315S, 1:2000; anti-pmTOR #2971S, 1:1000; anti-mTOR #2972S, 1:1000 (Cell Signaling); (2) Glutamate signaling: rabbit anti-Vesicular Glutamate Transporter 1, VGLUT1 #12331S, 1:500; anti-Postsynaptic Protein Density 95, PSD95 #3450S, 1:3000; anti-GluA1 #13185S, 1:2000 (Cell Signaling); (3) GABA signaling: rabbit anti- Vesicular GABA Transporter 1, VGAT #AB5062P, 1:2000 (Millipore); anti-gephyrin #AB5725, 1:500 (Millipore); anti-Glutamate Decarboxylase 1, GAD1 #5305S, 1:1000 (Cell Signaling). Appropriate secondary antibodies were used (1:5000, Vector Laboratories), under 1-2 h incubation, except for VGLUT1 that underwent overnight incubation. Bands were visualized with chemiluminescent reagent (Clarity Western ECL Substrate, Bio-Rad Laboratories, Inc., USA). Unsaturated and preserved bands were quantified using the Image Studio Lite software (v5.2, LICORbio, USA). GAPDH (#2118S rabbit, 1:5000, Cell Signaling) was used for loading control and normalization.

### Statistical analysis

Sample sizes were chosen based on previous experience with the tests employed and on power analyses (Cohen’s d power analysis, > 0.8 effect size). Results were analyzed by two-tailed paired or unpaired Student’s t-test, one- or two-way ANOVA as appropriate, followed by Dunnett’s multiple comparisons test, p ≤ 0.05. Results that did not follow a normal distribution were analyzed by Mann–Whitney U test or Kruskal–Wallis H test, p ≤ 0.05. Effect size was calculated using partial eta squared (η^2^_p_). The statistical tests used and the number of animals per group are listed in the figure and/or figure captions. Pre-established exclusion criteria included identified outliers (ROUT = 1%), mistargeted guide cannulas or lack of virus expression. All data sets were tested for homogeneity of variances using Levene’s test. All analyses were performed with SPSS Statistics (IBM, Version 30.0). Each experiment was independently replicated at least twice.

## RESULTS

### BSM facilitates motivational, hedonic and active coping responses without inducing drug-context association

We previously demonstrated that BSM produces fast (1 h) and sustained (at least up to 48 h) antidepressant-like effects in male mice, including increased natural reward-seeking behavior, motivation and active coping across a battery of behavioral tests relevant to antidepressant efficacy (27). Here, to determine whether these effects depend on the specific behavioral assay or on the order of tests within the behavioral battery, we employed a distinct experimental timeline. Male mice received BSM (10 mg/kg, i.p.) and were tested for (1) rapid effects in the FST and OFT on day 0, (2) sustained effects in the SuST and FUST on day 1, and TST on day 2 (Fig. 1A). For comparison, the prototype rapid-antidepressant ketamine (10 mg/kg) was used as a positive control.

Like ketamine, BSM reduced immobility time in the FST 1 h post-administration compared to the vehicle-control group (Fig. 1B). To rule out confounding effects of drug-induced acute locomotor stimulation, previously reported for ketamine within the first 20-30 min post-injection (31, 58), mice were exposed to the OFT after the FST (Fig. S1A). BSM, similar to ketamine, diminished passive coping behavior (Fig. 1B) without altering locomotor activity at this timepoint (Fig. S1A-C). At 24 h post-administration, both BSM and ketamine increased self-grooming time in the SuST, and enhanced female urine sniffing during the FUST (Fig. 1C-D), with no significant differences observed during water sniffing (Fig. S1D). At 48 h post-treatment, BSM also decreased immobility time in the TST, consistent with ketamine’s effects (Fig. 1E).

In rodents, α5-GABA_A_R have a restricted distribution in the PFC and hippocampus (21, 36). Particularly, the mPFC is consistently implicated in MDD, in stress-induced disturbances in rodents, and in the mechanisms underlying rapid antidepressant actions (19, 59). To characterize the effects of BSM directly microinjected in the mPFC, we performed a dose-response curve and assessed behavioral outcomes using the same battery of tests shown in Fig. 1A (Fig. 1F-G). BSM reduced the immobility time in the FST 30 min post-microinjection compared to the vehicle-control group (Fig. 1H) at all doses tested (3, 10 and 30 ng), with no alterations in locomotor activity (Fig. S1E-F). Twenty-four hours after intra-mPFC microinjection, BSM increased self-grooming time in the SuST (10 ng, Fig. 1I), and the duration of female urine sniffing in the FUST (3 and 30 ng), while no effect was found in the water sniffing (Fig. 1J and S1G). Forty-eight hours post-microinjection, BSM (10 and 30 ng) decreased immobility time in the TST (Fig. 1K). Overall, our results suggest that both the rapid (1 h) and sustained (24 to 48 h) antidepressant-like effects of BSM are driven, at least in part, by the mPFC, and that all tested doses are effective depending on the timepoint or behavioral assay employed.

Despite its clinical use as a fast-acting antidepressant, ketamine raises major concerns regarding addiction as a potential adverse effect. In preclinical research, sub-anesthetic doses of ketamine were demonstrated to promote CPP and hyperlocomotion (58, 60) as well as to elicit robust dopamine transients in the *nucleus accumbens* (61), alike long-stablished addictive substances. Therefore, to further assess abuse liability of BSM in comparison to ketamine, we evaluated drug-induced CPP and locomotor stimulant effects in the CPP paradigm (Fig. 2A). In contrast to ketamine, BSM (10 mg/kg) did not increase distance moved or velocity during the first 20 min following drug administration (conditioning day 1, Fig. 2B). In addition, unlike ketamine, BSM did not induce CPP, as indicated by unchanged time spent in the drug-paired compartment during the post-conditioning test compared with either the unpaired compartment or baseline preference (Fig. 2C; S1H). Here, we selected a 20 mg/kg dose of ketamine as positive control to ensure robust CPP induction under our experimental conditions, although ketamine-induced CPP has also been reported at doses ranging from 3 to 10 mg/kg (62). To model stress-induced impairments in adaptive reward processing, we next administered chronic CORT in the drinking water for 21 days and assessed CPP expression when the least-preferred arm of a T-maze was paired with sucrose, a natural reward with hedonic value (Fig. 2D). Under these conditions, both ketamine and BSM reversed CORT-induced deficits in sucrose-paired CPP expression (Fig. 2E). Together, these findings indicate that BSM selectively restores adaptive reward processing disrupted by stress while lacking intrinsic reinforcing properties, thereby dissociating antidepressant-like efficacy from abuse liability.

**Figure 2.**
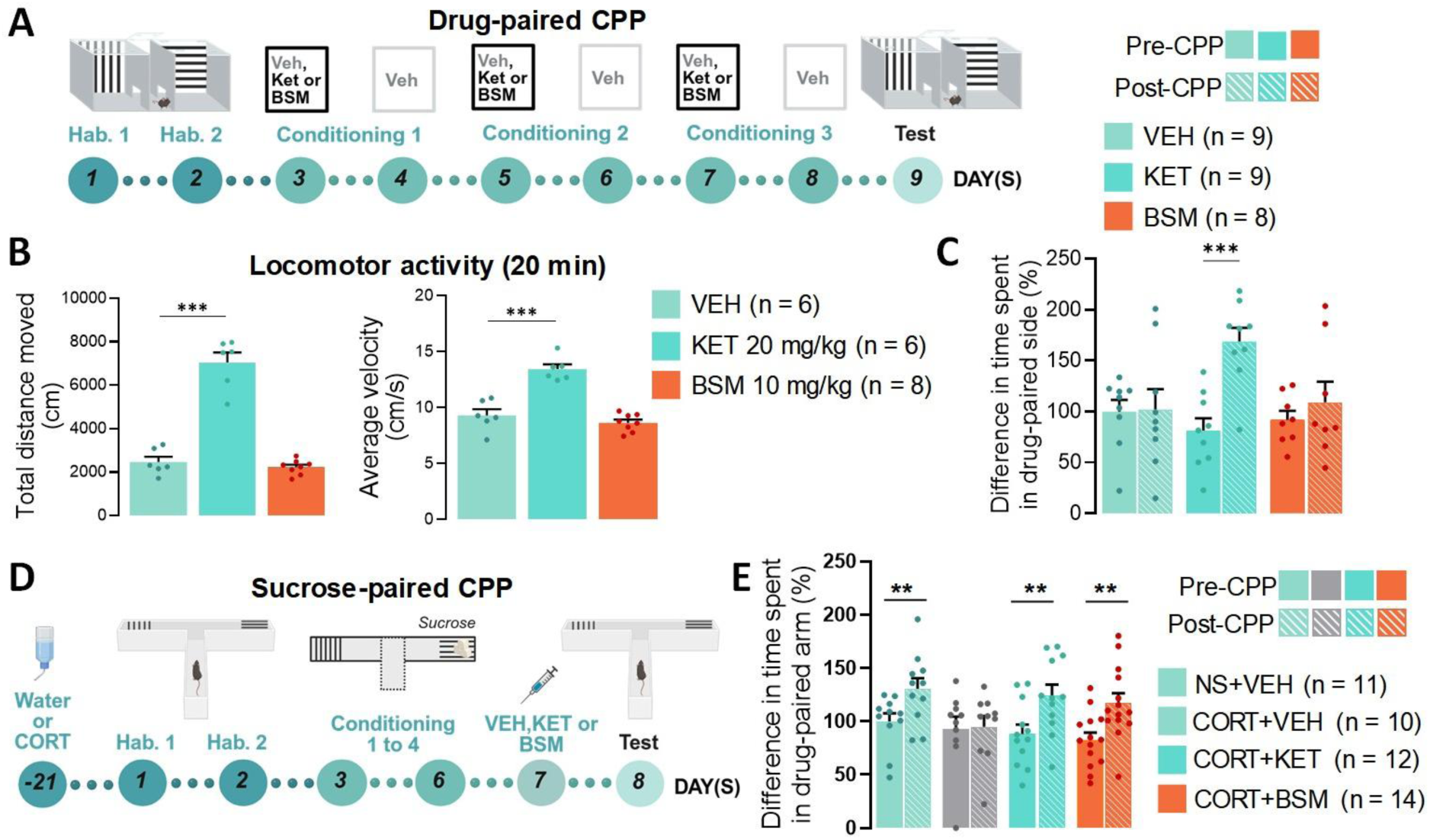
Basmisanil (BSM) facilitates sucrose- but not drug-paired conditioned place preference (CPP). **(A)** Schematic of the drug-paired CPP paradigm. Mice were habituated to the apparatus for two days (hab. 1–2) to establish baseline compartment preference (hab. 2, pre-CPP), then injected with vehicle (VEH), ketamine (KET, 20 mg/kg), or BSM (10 mg/kg) during six days of conditioning to one compartment, followed by a CPP test. **(B)** Average total distance traveled (H(2) = 12.11, P = 0.002) and velocity (F(2,17) = 38.26, P < 0.001) during the first drug–context pairing day. **(C)** Difference in time spent between the drug-paired and unpaired compartments within each test session (pre- and post-CPP), normalized to the VEH pre-CPP session (%; VEH: t(8) = 0.10, P = 0.92; KET: t(8) = 6.36, P < 0.001; BSM: t(7) = 1.05, P = 0.33). **(D)** Schematic of the sucrose-paired CPP paradigm. Mice received corticosterone (CORT) in the drinking water (0.1 mg/mL) for 21 days, were habituated to a T-maze for two days (hab. 1–2) to establish baseline arm preference (hab. 2, pre-CPP), and conditioned to sucrose in the least-preferred arm for four days (a.m./p.m.). Mice then received VEH, KET (10 mg/kg), or BSM (10 mg/kg), and CPP expression was assessed 24 h later. **(E)** Difference in time spent between the sucrose-paired arm and unpaired arms within each test session (pre- and post-CPP), normalized to the VEH pre-CPP session (%; non-stressed, NS+VEH: t(10) = 3.43, P = 0.0065; CORT+VEH: t(9) = 0.14, P = 0.89; CORT+KET: t(11) = 3.48, P = 0.0052; CORT+BSM: t(13) = 3.19, P = 0.0072). Data are shown as mean ± SEM. P values were calculated using independent-samples Kruskal-Wallis test, one-way ANOVA with Dunnett’s multiple-comparisons test or two-sided paired Student’s *t*-test. **P ≤ 0.01; ***P ≤ 0.001. Created with BioRender.

### BSM rapidly activates the mPFC and hippocampus, engaging mTOR signaling and enhancing glutamatergic and GABAergic synaptic function

Ketamine rapidly induces a transient glutamatergic burst in the mPFC, accompanied by GABAergic adaptations, resulting in activation of mPFC and hippocampal circuits and engagement of plasticity-related mechanisms (63–65). To examine whether BSM would change neuronal activity in target brain regions alike ketamine, we performed immunolabeling for c-Fos in the mPFC (PL and IL) and the hippocampus (dHC and vHC). Mice were euthanized 90 min after the injection of vehicle, BSM (10 mg/kg) or ketamine (10 mg/kg), and the number of c-Fos^+^ cells in each brain subregion was quantified per area (Fig. 3A). BSM, like ketamine, increased c-Fos expression in the PL and IL subregions of the mPFC (Fig. 3B and 3D), as well as in both the dHC and vHC, with the latter showing an approximately twofold elevation compared with vehicle-treated controls (Fig. 3C and 3E). Although ketamine increased the number of c-Fos^+^ cells in the vHC, c-Fos expression in the dHPC was comparable to that in vehicle-control mice (Fig. 3C). Representative hippocampal images indicate prominent BSM-induced c-Fos expression in CA1 subregions (Fig. 3E), consistent with previous reports indicating a high abundance of α5-GABA_A_R in this area (Sperk et al., 1997). Cells expressing dopamine D1 or D2 receptors comprise two major classes of excitatory pyramidal neurons in the PFC, typically referred to as Drd1- and Drd2-expressing neurons based on their gene identity (66). Specifically, D1R signaling mediates excitatory dopaminergic modulation across both superficial and deep cortical layers, with D1Rs highly enriched in pyramidal neurons of layers 5 and 6 (67) and required for rapid antidepressant-like responses (68). In contrast, within mPFC GABA interneurons subclasses, D1R expression is found predominantly in parvalbumin (PV) interneurons (66). Given evidence that the IL mPFC is preferentially engaged in rapid antidepressant-like responses relative to the PL region (63), we examined whether BSM activates D1R-expressing neurons in the IL (Fig. 3F-G). Ninety minutes after administration, both BSM and ketamine significantly increased c-Fos expression in D1R^+^ IL neurons compared with control mice (Fig. 3F). Next, to further characterize GABAergic recruitment, we conducted a double immunofluorescence for c-Fos and SST or PV in the IL. Drug treatments increased c-Fos expression selectively in SST^+^ interneurons, with no activation observed in PV^+^ cells (Fig. 3F). Altogether, we demonstrate that specific subpopulations of IL neurons, including DR1-expressing neurons, and SST-expressing interneurons, are engaged by both BSM and ketamine.

**Figure 3.**
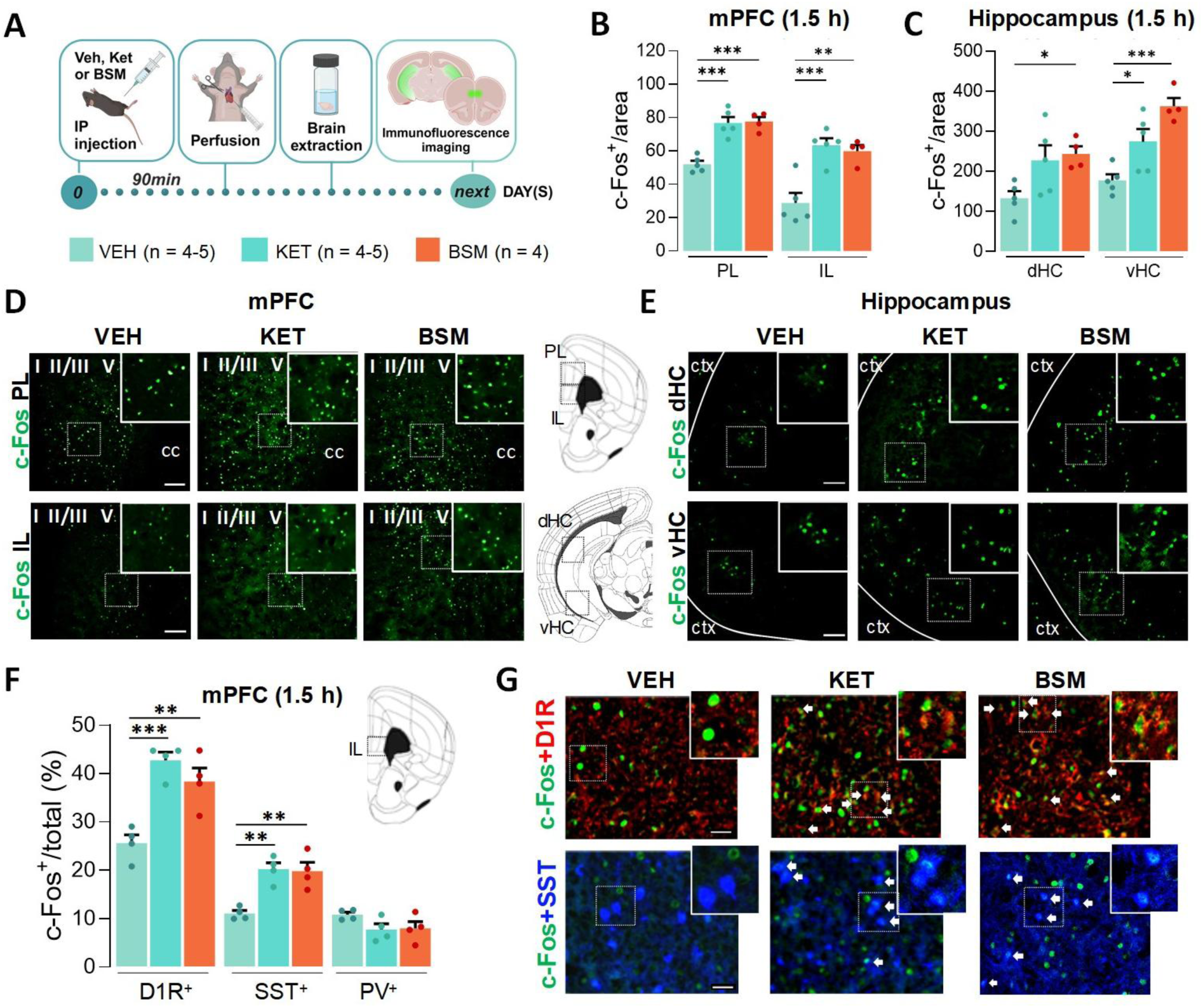
Basmisanil (BSM) induces expression of the activity marker c-Fos in mPFC D1R- or SST-expressing neurons and in the hippocampus. (**A**) Mice were injected with vehicle (VEH), ketamine (KET, 10 mg/kg, i.p.) or BSM (10 mg/kg, i.p.) and perfused 1.5 h post-treatment for c-Fos immunolabeling. (**B**) Quantification of the number of c-Fos^+^ cells per area in the prelimbic (PL) (F(2, 11) = 25.32; P < 0.001) and infralimbic (IL) mPFC (F(2, 11) = 15.74; P < 0.001), as well as (**C**) dorsal (dHC, F(2, 11) = 4.76; P=0.03) and ventral hippocampus (vHC, F(2, 11) = 14.35; P < 0.001). (**D**) Representative c-Fos immunofluorescence images in the PL (**top row**) and IL mPFC (**bottom row**; scale bar: 120 μm; magnification: ×10). (**E**) Representative c-Fos immunofluorescence images in the dHC (**top row**) and vHC (**bottom row**; scale bar: 100 μm; magnification, ×10). (**F**) Percentage of c-Fos^+^ cells that colocalize with D1R^+^, SST^+^ or PV^+^ neurons in the mPFC IL of VEH-, KET- and BSM-injected mice (D1R^+^: F(2, 9) = 17.37; P < 0.001. SST^+^: F(2, 9) = 14.54; P < 0.01. PV^+^: F(2, 9) = 2.34; P = 0.15). (**G**) Representative immunofluorescence images of IL mPFC cells from VEH, KET- or BSM-injected mice. White arrows indicate examples of colocalization (Magnification ×40). Data are plotted as mean ± SEM. P values were calculated using one-way ANOVA with Dunnett’s multiple-comparisons test. *P ≤ 0.05; **P ≤ 0.01; ***P ≤ 0.001. Created with BioRender.

Ketamine’s rapid antidepressant effects depend on neural plasticity (69) driven by mTOR-dependent synaptic signaling and the formation of new dendritic spines in the PFC (56). To investigate whether BSM engages similar cellular signaling pathways, we quantified synapse associated proteins in mPFC and hippocampal synaptoneurosomes at 1 and 24 h post-treatment (Fig. 4A). In the mPFC (Fig. 4B), 1h post-treatment, BSM induced a rapid increase in the phosphorylated (activated) forms of AKT, GSK3β, and mTOR, as well as in the levels of the AMPA glutamate receptor subunit GluA1. Conversely, BSM did not change phosphorylated levels of ERK 1/2.

**Figure 4.**
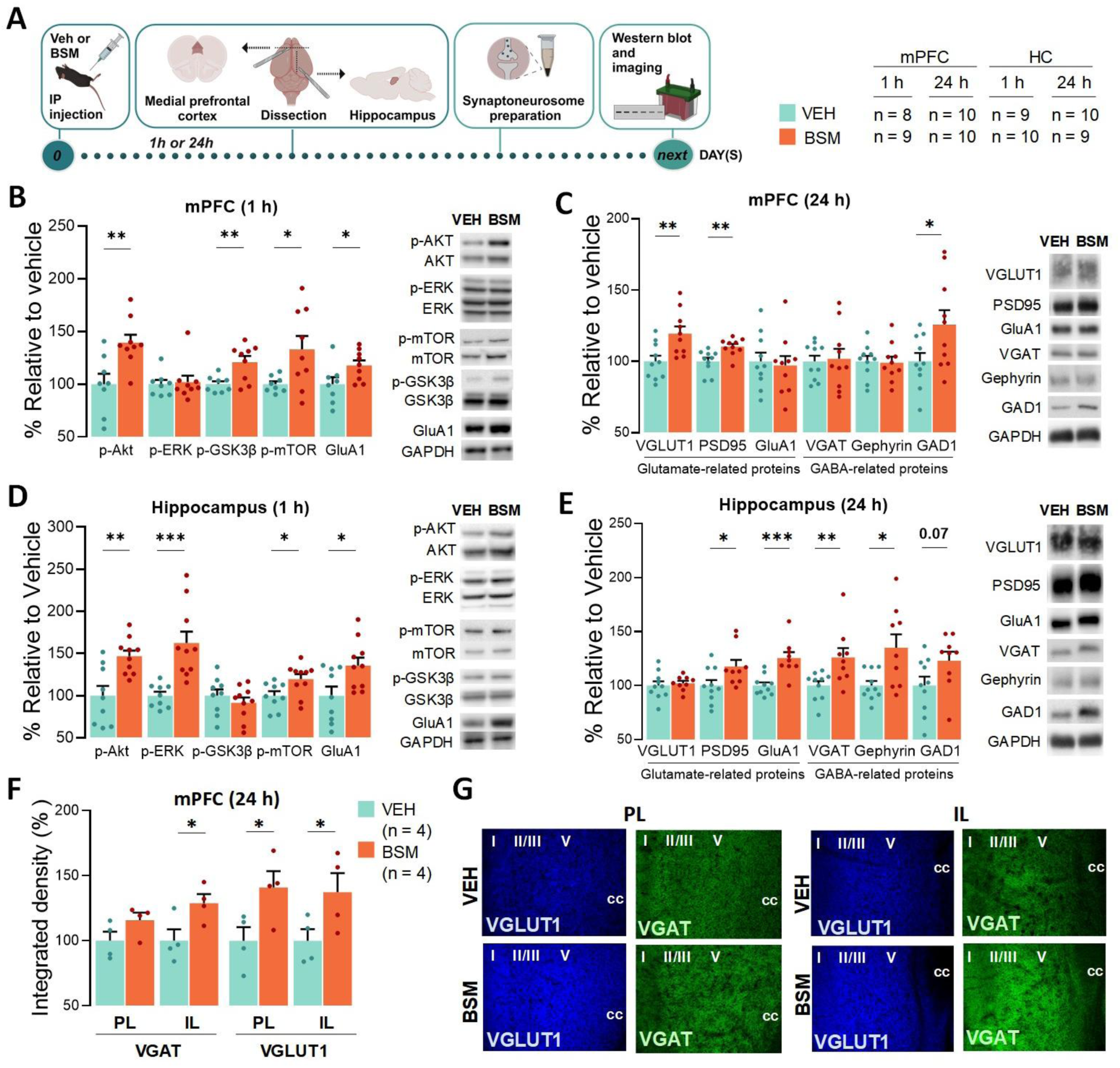
Basmisanil (BSM) activates the mTORC1 pathway and enhances synaptic proteins associated with both glutamatergic and GABAergic signaling in the mPFC and hippocampus (HC). (**A**) Mice were injected with either vehicle (VEH) or BSM (10 mg/kg, i.p.) and euthanized 1 h or 24 h post-treatment for Western Blotting analysis. After brain collection, mPFC and HC were dissected for subsequent synaptoneurosome preparation and protein quantification. (**B**) Quantification of immunoblotted signals in the mPFC from VEH- and BSM-injected mice 1 h post-treatment (p-AKT: t(15) = -3.23; P = 0.006. p-ERK: t(15) = -0.26; P = 0.80. p-GSK3β: t(15) = 3.29; P = 0.007. p-mTOR t(15) = 2.57; P=0.03, and GluA1: t(15) = -2.20; P = 0.04). (**C**) Quantification of immunoblotted signals in the mPFC from VEH- and BSM-injected mice 24 h post-treatment (VGLUT1: t(18) = -3.05; P = 0.007. PSD95: t(18) = -3.28; P = 0.004. GluA1: t(18) = 0.32; P = 0.75. VGAT: t(18) = -0.22; P = 0.83. Gephyrin: t(18) = 0.16; P = 0.88, and GAD1: t(18) = -2.20; P = 0.04. (**D**) Same as (B) but for hippocampal samples (p-AKT: t(17) = -3.60; P = 0.002. p-ERK: t(17) = -4.34; P = 0.001. p-GSK3β: t(17) = 0.87; P = 0.40. p-mTOR: t(17) = -2.44; P = 0.03, and GluA1: t(17) = -2.54; P = 0.02). (**E**) Same as (C) but for hippocampal samples (VGLUT1: t(17) = -0.46; P = 0.65. PSD95: t(17) = -2.21; P = 0.04. GluA1: t(17) = -4.30; P < 0.001. VGAT: t(17) = -2.78; P = 0.01. Gephyrin: t(17) = -2.67; P = 0.02, and GAD1: t(17) = -1.96; P = 0.07). All values were normalized with the respective non-phosphorylated protein or GAPDH, and the average value for vehicle mice was set in 100% (bar graphs start at 50%). (**F**) Percentage of integrated density of VGAT expression in the PL (t6 = 1.72, P = 0.14) and IL (t6 = 2.58, P = 0.04) and VGLUT1 expression in the PL (t6= 2.50, p = 0.04) and IL (t6= 2.19, P = 0.05) 24 h post-administration of VEH or BSM. (**G**) Representative VGAT and VGLUT1 immunofluorescence images (Magnification 20×). Data are plotted as mean ± SEM. P values were calculated using two-sided unpaired Student’s t test. *P ≤ 0.05; **P ≤ 0.01; ***P ≤ 0.001. Created with BioRender.

Activation of mTOR signaling has been linked to plasticity at both glutamatergic and GABAergic synapses (70). To assess sustained synaptic adaptations in glutamate and GABA signaling, we quantified neurotransmitter-related proteins in mPFC synaptoneurosomes 24 h post-treatment (Fig. 4C). Relevant to mPFC glutamatergic signaling, BSM administration increased levels of VGLUT1, which mediates glutamate loading into synaptic vesicles, and PSD95, a major scaffolding component of excitatory synapses (Fig. 4C). BSM did not produce significant changes in GluA1 at 24 h, indicating that the rapid elevation observed 1 h after administration is not maintained at later time points. For GABA signaling, BSM significantly increased levels of GAD1, the predominant enzyme isoform that catalyzes the synthesis of GABA from glutamate (Fig. 4C). No late changes were observed in the levels of VGAT or gephyrin, a major scaffolding protein at inhibitory postsynaptic densities, in mPFC homogenates. To further investigate whether putative alterations in GABA-related proteins would be restricted to mPFC subregions (PL or IL) rather than the whole-tissue homogenate, we measured VGAT and VGLUT1 expression in mPFC slices, and found that BSM-induced increase in VGAT levels is restricted to the IL but not PL (Fig. 4F-G). Conversely, VGLUT1 expression was increased in both mPFC subregions, indicating that while glutamatergic changes are more widespread, BSM might enhance GABAergic signaling in a more circuit-specific manner.

In the hippocampus, BSM significantly increased the phosphorylated forms of AKT, ERK, and mTOR, as well as GluA1 levels, 1 h post-administration, with no effect on p-GSK3β expression (Fig. 4D). At 24h, we also found increased levels of PSD95 and GluA1, with no change in VGLUT1 (Fig. 4E). VGAT and gephyrin expression were also elevated, while GAD1 showed a trend towards significance (Fig. 4E). These results indicate that BSM-induced facilitation of synaptic plasticity is also present in the hippocampus, a region with high α5-GABA_A_R expression, warranting future studies.

### BSM promotes behavioral recovery following chronic stress exposure

The CUS paradigm models maladaptive behaviors in rodents that recapitulate key features of stress-related states, including impairments in motivation, cognition, and sociability (42, 71). Here, mice were exposed to CUS for 21 days and underwent treatment, followed by behavioral testing in the NOR test (1h – test and 24 h – retest), FUST, SuST, and FST (Fig. 5A). At both 1 and 24 h post-treatment, CUS decreased the discrimination index in vehicle-treated mice relative to non-stressed controls, indicating impaired recognition of the familiar object (Fig. 5B-C; Fig. S2A-D). Notably, these changes were reversed by treatment with either BSM or ketamine (Fig 5B-C). CUS significantly reduced sniffing time to female urine during the FUST (26 h post-treatment, Fig. 5D) and self-grooming in the SuST (48 h, Fig. 5E), as well as increased immobility time in the FST (50 h, Fig. 5F) – effects that were significantly reversed by BSM or ketamine. No changes were observed in the water sniffing time during the FUST (Fig. S2E).

**Figure 5.**
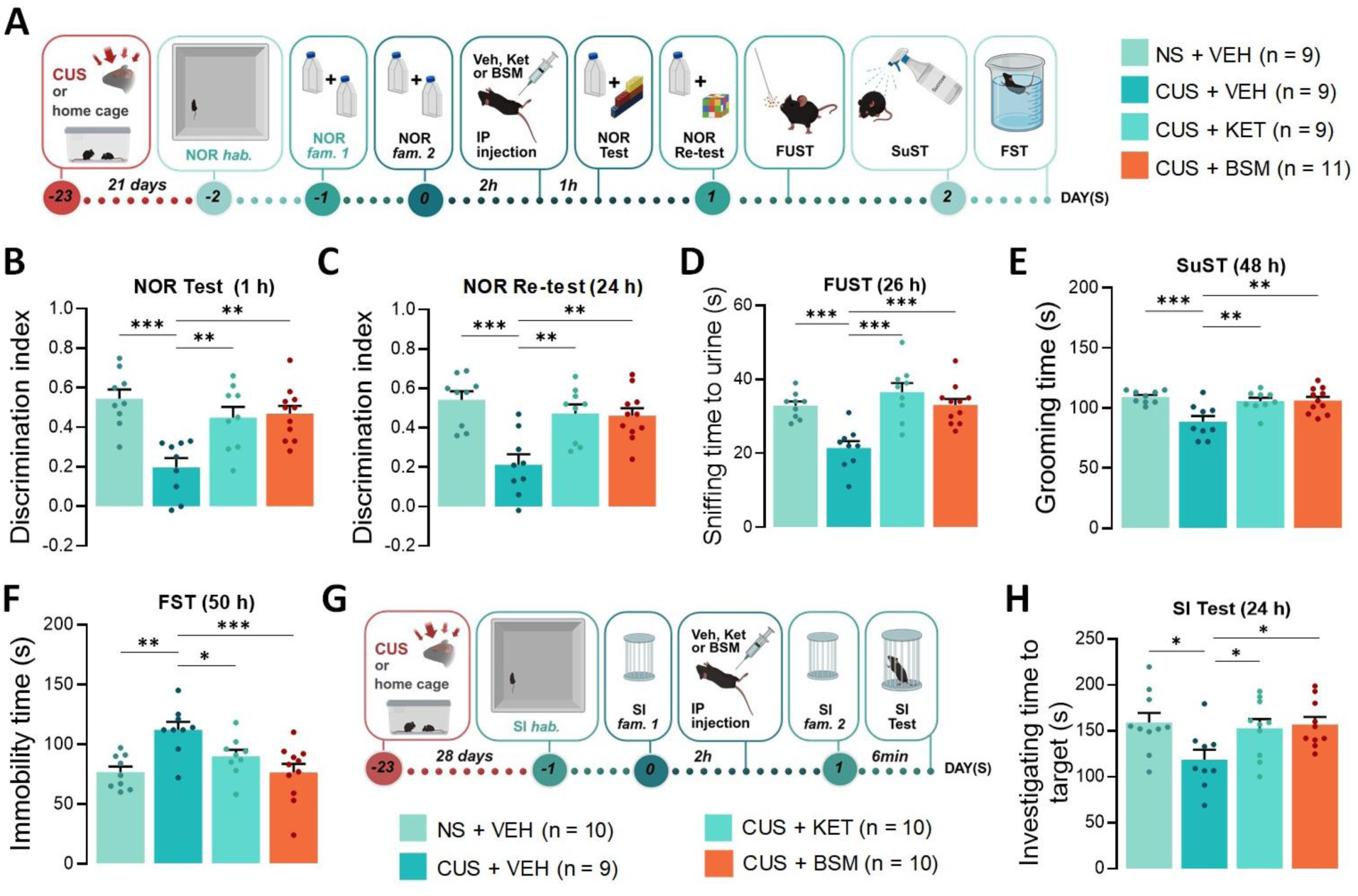
Basmisanil (BSM) reverses maladaptive behaviors induced by chronic unpredictable stress (CUS) in male mice. (**A**) Mice underwent 21 days of CUS followed by behavioral tests performed 1, 24, 26, 48 and 50 h after a single injection of vehicle (VEH), ketamine (KET 10 mg/kg, i.p.) or BSM (10 mg/kg, i.p.). Non-stressed (NS) mice remained in the home cage and were injected with VEH. (**B**) Average discrimination index during the novel object recognition (NOR) test 1 h (F(3,34) = 9.82; P < 0.001), or (**C**) 24 h post-treatment (F(3,34) = 10; P < 0.001). (**D**) Average time spent sniffing female urine during the female urine sniffing test (FUST, F(3,34) = 12.30, P < 0.001), (**E**) self-grooming during the sucrose splash test (SuST, F(3,34) = 7.60, P < 0.001), or (**F**) immobile during the forced swimming test (FST, F(3,34) = 7.13, P < 0.001). (**G**) Mice underwent 28 days of CUS, with social isolation during the final 21 days, followed by the social interaction (SI) test performed 24 h after a single i.p. injection of VEH, KET (10 mg/kg) or BSM (10 mg/kg). Non-stressed mice remained group-housed in the home cage and were injected with VEH. (**H**) Average time spent investigating an unfamiliar social target (F(3,35) = 3.53, P = 0.02). Data are plotted as mean ± SEM. P values were calculated using one-way ANOVA with Dunnett’s multiple-comparisons test. *P ≤ 0.05; **P ≤ 0.01; ***P ≤ 0.001. Created with BioRender.

Social isolation is an adverse environmental factor associated with the development of stress-related maladaptive states (72). A single dose of racemic ketamine or (R)-ketamine restores anhedonia-like behaviors and social memory deficits in male rodents socially isolated during adulthood or post-weaning (51, 73). In this context, we examined whether BSM could rescue social approach impaired by CUS in combination with chronic social isolation (Fig. 5G). Stressed mice exhibited significantly less time investigating the social target during the test, and this deficit was restored by BSM or ketamine 24 h post-treatment (Fig. 5H). The time investigating the empty enclosure was comparable among all groups (Fig. S2G). Together, these results indicate that BSM promotes ketamine-like effects in counteracting CUS-induced maladaptive changes in cognitive function, reward-seeking, self-care, coping and social behaviors from 1 to 50 h post-treatment.

### Rapid recruitment of mPFC CaMKIIα^+^ neurons mediates BSM’s behavioral effects

It is unknown whether activity of α5-GABA_A_R in excitatory neurons, interneurons or both is required to mediate the behavioral effects of α5-NAMs. Given that the majority of cortical α5-GABA_A_Rs are in pyramidal cells of human (39.7%) and mouse (54.14%) frontal cortex (74), we hypothesized that BSM promotes excitatory disinhibition through direct negative allosteric modulation of α5-GABA_A_Rs in pyramidal neurons. To test this, we performed chemogenetic inhibition of mPFC CaMKIIα^+^ neurons by microinjecting a Cre-dependent hM4DGi virus into the mPFC of CaMKIIα^Cre+^ or WT mice (Fig. 6A). After surgical recovery, animals received i.p. injection of CNO (1 mg/kg), followed by either vehicle or BSM (10 mg/kg) 20 min later (Fig. 6A). CNO has a short half-life in mice (∼1 h) (40), thus this timeframe enables effective silencing of glutamatergic neurons during the initial phase of BSM’s actions, with potential consequences for subsequent plasticity-related processes. Behavioral testing began 1 h after treatments, and mice were tested for up to 48 h.

**Figure 6.**
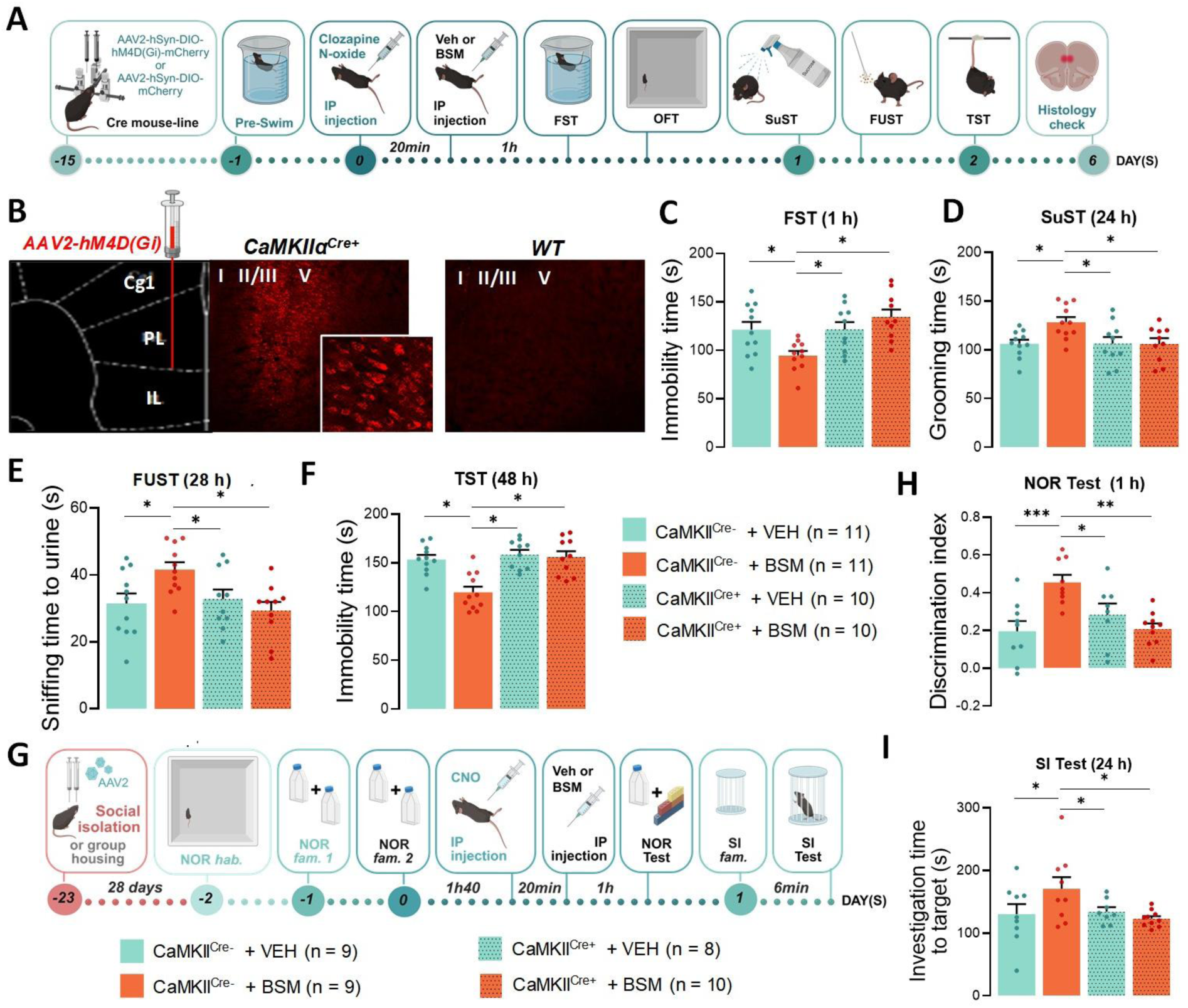
Chemogenetic inhibition of mPFC CaMKII^+^ neurons blocks both the rapid and sustained behavioral actions of Basmisanil (BSM). (**A**) Timeline for surgery, treatments, and behavioral testing. **(B)** Representative images of the mPFC from CaMKII^Cre+^ and WT (Cre-negative littermate) mice that received bilateral infusions of hM4DGi virus (Magnification: ×10; inset: ×20+zoom). Behavioral tests were performed 1, 24, 28 and 48 h after injection of either vehicle (VEH) or BSM (10 mg/kg, i.p.). All groups were injected with CNO (1 mg/kg) 20 min prior to VEH or BSM. (**C**) Average time spent immobile during the forced swimming test (FST, F_int_ (1, 38) = 7.87, P=0.008; η^2^_p_ effect size = 0.17; F_gen_ (1, 38) = 7.98, P=0.007; F_treat_ (1, 38) = 0.92, P = 0.34), (**D**) self-grooming during the sucrose splash test (SuST, F_int_ (1, 38) = 4.14; P = 0.05; η^2^_p_ effect size = 0.10; F_gen_ (1, 38) = 3.98, P = 0.05; F_treat_ (1, 38) = 3.92, P = 0.05), (**E**) sniffing female urine during the female urine sniffing test (FUST, F_int_ (1, 38) = 6.50; P = 0.01; η^2^_p_ effect size = 0.15; F_gen_ (1, 38) = 4.19, P = 0.05; F_treat_ (1, 38) = 1.55, P = 0.22), and (**F**) immobile during the tail suspension test (TST; F_int_ (1, 38) = 8.45; P = 0.006; η^2^_p_ effect size = 0.18; F_gen_ (1, 38) = 15.01, P < 0.001; F_treat_ (1, 38) = 11.66, P = 0.002). (**G**) hM4D(Gi)- CaMKII^Cre+^ and -WT underwent the novel object recognition (NOR) test and social interaction (SI) test 1 and 24 h, respectively, after a single injection of VEH or BSM (10 mg/kg, i.p.). All groups were injected with CNO 20 min before VEH or BSM. (**H**) Average discrimination index of a novel object during the NOR test (F_int_ (1, 31) = 21.51; P < 0.001; η^2^_p_ effect size = 0.43; F_gen_ (1, 31) = 1.11, P = 0.30; F_treat_ (1,31) = 1.24, P = 0.27), and (**I**) time spent investigating an unfamiliar social target during the SI test (F_int_ (1,31) = 6.03; P = 0.02; η^2^_p_ effect size = 0.16; F_gen_ (1,31) = 1.96, P = 0.17; F_treat_ (1,31) = 2.39, P = 0.13). Data are plotted as mean ± SEM. P values were calculated using two-way ANOVA with Dunnett’s multiple-comparisons test. *P ≤ 0.05; **P ≤ 0.01; ***P ≤ 0.001. Figure created with BioRender.

Fluorescence images revealed high hM4D(Gi)-mCherry^+^ density in the intermediate and deep layers of mPFC in CaMKIIα^Cre+^ mice (Fig. 6B). As expected, BSM decreased immobility time in the FST (1 h), increased grooming time in the SuST (24 h) and sniffing time of female urine in the FUST (28 h), and reduced immobility in the TST (48 h) in WT mice. All of these effects were prevented by chemogenetic inhibition of CaMKIIα^+^ neurons across all testing timepoints in CaMKIIα^Cre+^ mice compared to WT littermates. There were no significant changes on locomotor activity during the OFT or sniffing time toward water during the FUST (Fig. S3A-C). Noteworthy, pre-treatment with a low dose of CNO *per se* did not change any behavior in CaMKIIα^Cre+^ mice treated with vehicle. Altogether, these findings demonstrate that activity at mPFC glutamatergic synapses is required for both the rapid and sustained effects of BSM on motivated behavior, reward seeking, and coping strategies.

We also investigated cognitive and social domains by testing independent cohorts of CaMKIIα^Cre+^ mice and WT littermates in the NOR test (1 h) followed by SI test (24 h). After intra-mPFC AAV-hM4D(Gi) infusion, mice remained single housed during the entire period of recovery and behavioral testing (28 days, Fig. 6G), since chronic social isolation induces cognitive impairment, and social behavior deficits in rodents (75). As suggested for ketamine (13), single housing may increase baseline stress levels, thereby permitting more robust behavioral responses to rapid-acting antidepressants. Here, BSM increased the discrimination index during the NOR test (1 h) and time spent investigating a social target during the SI test (24 h) in WT mice, and these effects were prevented by chemogenetic inhibition of CaMKIIα^+^ neurons. During familiarization, investigation times for the familiar objects in the NOR test and the empty enclosure in the SI test remained unchanged (Fig. S3F). Additionally, CNO treatment alone did not affect outcomes in any of the behavioral assays. Together, these results suggest that BSM triggers rapid CaMKIIα^+^ neuronal activity in the mPFC, facilitating object discrimination memory and social approach in male mice.

### mPFC GAD1^+^ interneurons sustain BSM’s behavioral effects

Although the *GABRA5* gene is expressed in frontal cortical GABAergic interneurons to a much lesser extent (74), rapid glutamatergic disinhibition may nonetheless drive secondary GABAergic adaptations that are relevant to antidepressant efficacy (47). Therefore, we also investigated whether α5-GABA_A_R in GABA interneurons are required for the behavioral responses induced by BSM. For comparison, we repeated the same experiments described above in Gad^Cre^ mice to chemogenetically inhibit GABAergic interneurons 20 min before BSM or vehicle treatment (Fig. 7A). In WT mice, BSM decreased immobility in the FST (1 h) and increased grooming time in the SuST (24 h) (Fig. 7B-C). These effects were attenuated but not blocked by CNO treatment in GAD1^Cre+^ mice. In contrast, at later timepoints, inhibition of mPFC GABA interneurons completely reversed BSM’s effects during the FUST (Fig. 7D) and TST (Fig. 7E) 28 h and 48 h post-treatment, respectively. No changes were observed in locomotor activity in the OFT or in sniffing time toward water during the FUST (Fig. S3G-I). Because the impact of chemogenetic silencing in CaMKIIα^Cre+^ and GAD1^Cre+^ neurons on blocking BSM’s effect differed in both magnitude and time course, η^2^_p_ effect size was used to compare the estimated interaction effect sizes between experiments combining drug treatment and chemogenetic silencing. For the experiment involving CaMKIIα^+^ neuron silencing (interaction treatment*genotype), η^2^_p_ effect size in the FST (1 h), SuST (24 h), FUST (28 h) and TST (48 h) was 0.17, 0.10, 0.15 and 0.18, respectively. For Gad^Cre+^ interneuron silencing, η^2^_p_ effect sizes were 0.10, 0.09, 0.18 and 0.19, respectively. From a longitudinal perspective, these results suggest that the effect magnitude is initially greater following CaMKIIα^+^ neuron inhibition, beginning as early as 1 h post-treatment, whereas the impact of GAD1^Cre+^ neuron inhibition progressively increases over time.

**Figure 7.**
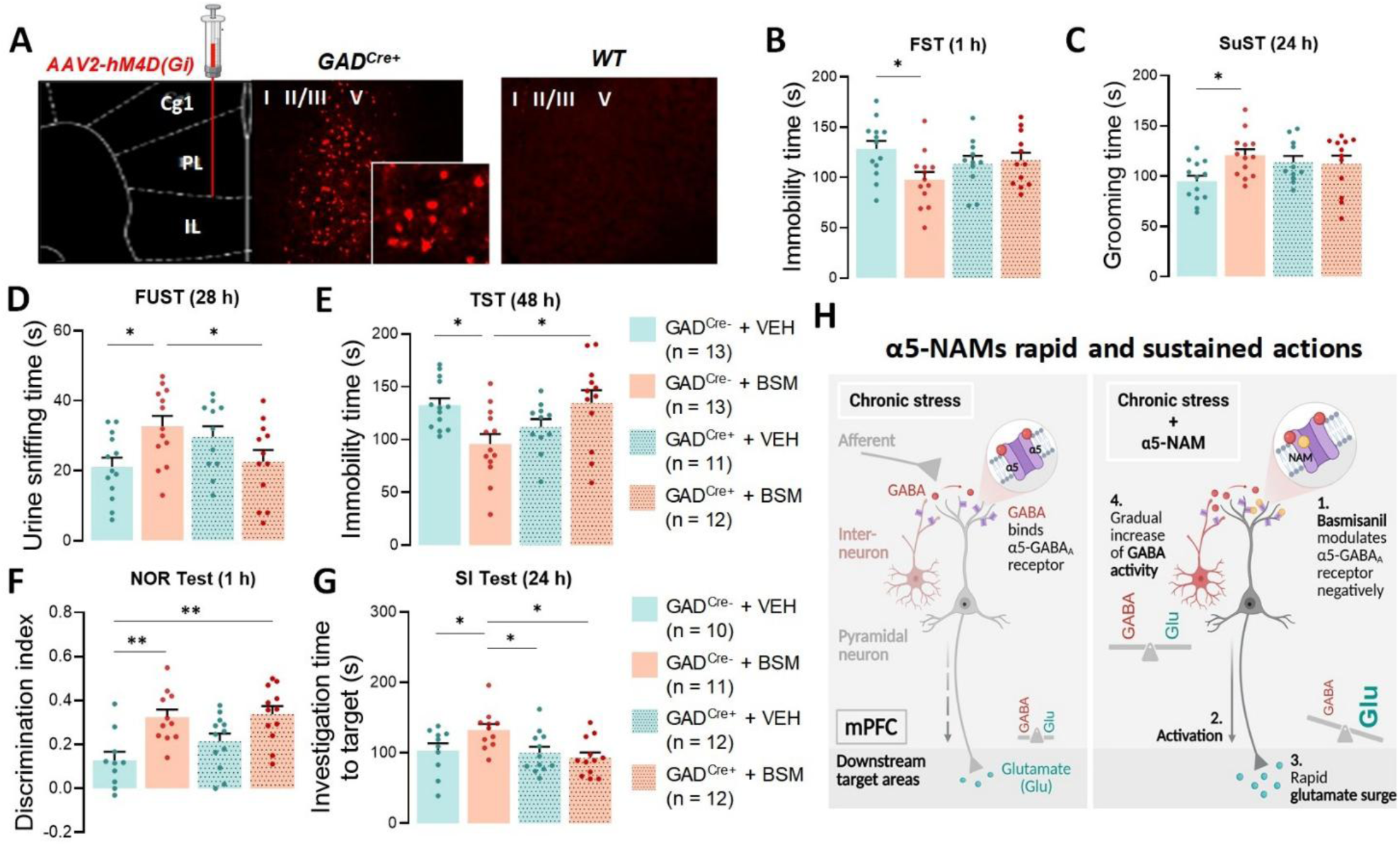
Chemogenetic inhibition of mPFC GAD^+^ neurons blocks the sustained but not rapid behavioral actions of Basmisanil (BSM). Timeline for surgery, treatments, and behavioral testing are as shown in figure 6A, but performed in GAD^Cre+^ mice and WT littermates. **(A)** Representative images of the mPFC from GAD^Cre+^ and WT (Cre-negative littermate) mice that received bilateral infusions of hM4DGi virus (Magnification: ×10; inset: ×20+zoom). Behavioral tests were performed 1, 24, 28 and 48 h after injection of VEH or BSM (10 mg/kg, i.p.). All groups were injected with CNO (1 mg/kg) 20 min prior VEH or BSM. (**B**) Average time spent immobile during forced swimming test (FST, F_int_ (1, 45) = 4.80, P = 0.03; η^2^ effect size = 0.10; F_gen_ (1, 45) = 0.08, P=0.77; F (1, 45) = 3.16, P=0.08), (**C**) self-grooming during the sucrose splash test (SuST, F_int_ (1, 45) = 4.27; P = 0.04; η^2^_p_ effect size = 0.09; F_gen_ (1, 45) = 0.63, P=0.43; F_treat_ (1, 45) = 3.43, P=0.07), (**D**) sniffing female urine during the female urine sniffing test (FUST, F_int_ (1, 45) = 9.80; P = 0.003; η^2^ effect size = 0.18; F_gen_ (1, 45) = 0.07, P=0.80; F_treat_ (1, 45) = 0.58, P=0.45), and (**E**) immobile during the tail suspension test (TST, F_int_ (1, 45) = 10.77; P = 0.002; η^2^ effect size = 0.19; F_gen_ (1, 45) = 1.04, P=0.31; F (1, 45) = 0.61, P=0.44). **(F-G)** hM4D(Gi)-CaMKII^Cre+^ and -WT underwent the novel object recognition (NOR) test and social interaction (SI) test 1 and 24 h, respectively, after a single injection of VEH or BSM (10 mg/kg, i.p.). All groups were injected with CNO 20 min prior VEH or BSM. (**F**) Average discrimination index of a novel object in the NOR test (F_int_ (1,41) = 0.99; P = 0.32, η^2^ effect size = 0.02; F_gen_ (1,41) = 1.89, P = 0.18; F_treat_ (1,41) = 18.71, P < 0.001), and (**G**) time spent investigating an unfamiliar social target in the SI test (F_int_ (1,41) = 4.43; P = 0.04, η^2^ effect size = 0.10; F_gen_ (1,41) = 6.08, P = 0.02; F_treat_ (1,41) = 1.63, P = 0.21). Data are plotted as mean ± SEM. P values were calculated using two-way ANOVA with Dunnett’s multiple-comparisons test. *P ≤ 0.05; **P ≤ 0.01. (**H**) Working model for the rapid and sustained actions of α5-NAMs in the mPFC. **Left panel:** Under chronic stress, mPFC circuit function is disrupted, resulting in altered excitatory–inhibitory (E/I) balance within pyramidal neuron networks. Given that intact mPFC function relies on precise E/I regulation to support higher-order processes such as emotional regulation, working memory, and decision-making, these circuit-level disturbances lead to degraded information coding and reduced transmission to downstream regions. **Right panel:** Acute α5-NAM administration produces a rapid reduction in α5-GABA_A_R-mediated inhibitory tone on mPFC pyramidal neurons, where α5 subunits are highly expressed, resulting in prefrontal disinhibition and fast (∼1 h) behavioral improvements. This initial increase in excitatory drive is proposed to engage downstream adaptive processes, including enhanced GABAergic activity and synaptic plasticity within both glutamatergic and inhibitory neurons, which persist for days. Together, these mechanisms restore mPFC E/I balance and support sustained recovery of emotional and cognitive function. Created with BioRender.

We next tested independent cohorts of GAD1^Cre+^ mice in the NOR test followed by the SI test 1 and 24 h post-administration of BSM. Notably, these assays strongly depend on intact cortical function (75–77) and are therefore particularly informative at the circuit level. Unlike CaMKIIα^Cre+^ mice (Fig. 6H), chemogenetic silencing of mPFC GABAergic interneurons in Gad1^Cre+^ mice did not reverse or attenuate BSM-induced enhancement of object recognition memory at 1 h post-treatment (Fig. 7F; Fig. S3J), but did reverse the BSM-induced increase in social interaction observed in the SI test 24 h later (Fig. 7G). Neither chemogenetic nor pharmacological manipulations were associated with changes in total object exploration in the NOR test or investigation time of empty enclosures in the SI test (Fig. S3K-L). Overall, our findings are consistent with our previous report for ketamine (13), suggesting that GABAergic adaptations following BSM administration are more critical for sustaining long-term behavioral effects than for mediating rapid responses.

## DISCUSSION

Here, we show that BSM promotes rapid and sustained antidepressant-like effects and alleviates stress-induced behaviors via mPFC molecular adaptations and time-dependent recruitment of excitatory and inhibitory circuits, phenotypes that parallel ketamine’s actions (14, 47). In addition to cortical involvement, our findings further indicate that the hippocampus also exhibits increases in neuronal activity and molecular plasticity in response to α5-GABA_A_R modulation, corroborating previous studies and highlighting this region as a promising target for future investigation (28, 33). Mechanistically, chemogenetic inhibition of mPFC pyramidal neurons abolishes the motivational, hedonic, active coping, procognitive and prosocial effects of BSM across both rapid and sustained timescales, whereas silencing of GABAergic interneurons reverses long-term behavioral improvements. These findings support a model in which, similar to ketamine (47), BSM drives an initial phase of rapid cortical excitation, impacting both short-term and subsequent long-term behavioral outcomes, and GABAergic adaptations that sustain these effects. Together, our data reveal molecular, cellular, and behavioral mechanisms underlying BSM’s actions, highlighting its potential as a novel, targeted antidepressant that restores cortical E/I balance and adaptive prefrontal function.

Due to their enriched spatial distribution in the mPFC and compelling preclinical evidence, α5-GABA_A_Rs have emerged as promising targets to novel antidepressant agents with the potential to minimize adverse side effects (24). Here, consistent with our previous findings, we demonstrate that either systemic or intra-mPFC administration of BSM produces fast (within 1 h) and long-lasting (investigated up to 48 h) behavioral responses in murine tasks without inducing ketamine’s typical hyperlocomotion or CPP. Notably, although BSM did not demonstrate efficacy in clinical trials in improving cognitive deficits associated with genetic neurodevelopmental conditions such as Down syndrome (35), its effects seem more sensitive to reversing cognitive and dendritic impairments induced by stress during early adulthood. This profile suggests that α5-NAMs hold greater promise as a rapid-acting antidepressant targeting stress-related synaptic dysfunction rather than as a therapeutic aimed at correcting developmental circuit abnormalities.

Distinct classes of GABAergic (such as SST and PV) and glutamatergic (including Drd1 and Drd2) neuronal populations play a crucial role in a variety of coordinated integrative brain functions, enabling the brain to adapt and process information through a precise balance between E/I signals (19, 78). In this context, our findings demonstrate that a single administration of BSM engages PFC circuitry similar to ketamine, eliciting robust c-Fos activation in both mPFC PL and IL subdivisions. Notably, this activation was especially prominent in D1R-expressing neurons within the IL, which accounted for approximately 40–50% of all c-Fos-positive cells and are described as predominantly pyramidal neurons (67). This observation aligns with prior evidence indicating that ketamine’s rapid and sustained antidepressant-like effects critically depend on the activation of Drd1-expressing neurons in the mPFC, and that selective stimulation of this population is sufficient to recapitulate ketamine-like behavioral outcomes (68).

Interestingly, we also identified that both BSM and ketamine induce neuronal activity in GABAergic SST interneurons within the IL, whereas no effect was observed in PV interneurons. Mechanistically, these effects are unlikely to reflect direct modulation of SST interneurons by BSM. Transcriptomic analyses indicate that α5-GABA_A_R subunits are highly enriched in excitatory pyramidal neurons (∼54%), expressed at much lower levels in PV interneurons (∼16%), and are largely absent from SST interneurons (74). Thus, as supported by our chemogenetic data, BSM likely acts initially on pyramidal neurons via α5-GABA_A_R modulation, enhancing excitatory output and triggering downstream circuit-level adaptations. One plausible consequence of this enhanced excitation is the recruitment of SST interneurons through feedback mechanisms that constrain dendritic excitability and prevent excessive glutamatergic drive (47, 79, 80). Activation of CaMKII^+^ glutamatergic neurons engages a positive feedback mechanism culminating in long-term potentiation of dendritic inhibition mediated by mPFC SST interneurons (81). In addition, SST interneurons exert inhibitory control over both pyramidal neurons and PV interneurons, and SST-to-PV connections within cortical microcircuits may account for the absence of detectable activity changes in PV cells observed here (82). Notably, both excitatory and inhibitory neurons have been shown to exhibit overlapping *Fos* gene expression between 60 and 120 min after stimulation (83), which may account for our detection of c-Fos induction in both populations of neurons at the 1.5 h timepoint. Importantly, although ketamine has been suggested to produce a rapid and transient blockade of SST interneurons (11), our findings are consistent with a biphasic model of GABAergic modulation, in which an initial reduction in interneuron activity (<30 min) is followed by a rebound beginning approximately 1 h after treatment (46). Nonetheless, further studies will be required to test causality and resolve functional heterogeneity within D1R^+^ and SST^+^ population ensembles following ketamine or BSM (13).

At the molecular level, extensive literature has shown deficits in synaptic markers, structural remodeling and synaptic plasticity in the PFC and hippocampus of MDD subjects and chronically stressed rodents, leading to reduced function of both glutamatergic and GABAergic signaling pathways (14, 24, 39, 43, 44, 84–86). Moreover, we have shown that rapid-acting agents, including ketamine and the acetylcholine muscarinic antagonist scopolamine, promote sustained enhancement of glutamate- and GABA-induced molecular plasticity (14, 44). As previously reported for these drugs and other rapid-acting therapeutics (14, 44, 56, 57, 87, 88), our results show that a single systemic administration of BSM increases synaptic activation of GluA1 and Akt- or Erk-mTOR signaling pathways in the mPFC and hippocampus. Following this initial response, we observed elevated levels of synaptic proteins associated to both glutamatergic and GABAergic signaling. Interestingly, VGAT expression was specifically increased in the IL but not PL mPFC, suggesting regional differences in inhibitory synaptic regulation that warrant further investigation. Indeed, both ketamine administration and optogenetic stimulation have been shown to preferentially recruit the IL, rather than the PL, to promote antidepressant-like effects (63). Consistent with our glutamatergic findings, the α5-GABA_A_Rs NAM MRK-016 increases cortical γ-frequency oscillations *in vivo*, an effect that is prevented by prior administration of the AMPA receptor antagonist NBQX, indicating activity-dependent synaptic potentiation provoked by the α5-NAM (31). In alignment, MRK-016 also restores AMPA-to-NMDA ratios in *ex vivo* hippocampal slices from chronically stressed mice (28). Although the primary focus of the present study is the mPFC, a recent report demonstrated that pharmacological enhancement of ERK activity transiently augments ketamine-induced synaptic potentiation at CA3-CA1 (89), supporting a role for BSM-mediated activation of ERK signaling in the hippocampus. Furthermore, we report that BSM induces expression of gephyrin in the hippocampus but not in the mPFC, indicating an upregulation of the scaffolding protein that anchors synaptic GABA_A_Rs. In hippocampal neurons, AMPA receptor activation increases the recruitment of α5-GABA_A_Rs to postsynaptic sites (23), where the α5 subunit is significantly colocalized with gephyrin (90). Thus, although α5-GABA_A_Rs are classically enriched at extrasynaptic locations, they can also be recruited to synaptic sites where they are necessary for dendritic outgrowth and spine maturation (90). Future studies examining the role of radixin, a protein that anchors α5-GABA_A_Rs to extrasynaptic membranes, may help clarify region-specific mechanisms engaged by BSM.

Complex human behavior is difficult to model in rodents, and we acknowledge the limitations, including relevance to depression and treatment response (91, 92). As discussed, the PFC undergo morphological and functional alterations in both MDD subjects and chronically stressed animals (93, 94), including disrupted projections and dysregulated downstream limbic activity (14, 39, 42, 44, 57, 94, 95), leading to maladaptive outcomes in motivational, cognitive and social behaviors. Most antidepressants have not been developed and/or evaluated for their ability to directly and independently ameliorate cognitive deficits (96). Notably, early findings with global α5 gene *knockout* or reduced α5-GABA_A_R expression in the hippocampal formation showed that this receptor subtype is involved in the physiology of learning and memory (20, 97). Here, we show that, in addition to completely reversing CUS-induced impairments in motivational, reward-seeking and coping behaviors in a fast and prolonged manner (up to 50 h post-injection), BSM also reverses deficits in object recognition memory, with this procognitive effect being both acute and long-lasting. Consistent with these findings, BSM improves spatial learning impairment in rats and executive function in cynomolgus macaques in hippocampal- and prefrontal cortex-dependent tasks, respectively (36). Our study further revealed a similar rescue of CUS-induced memory deficits by ketamine, corroborating previous reports showing enhanced NOR performance following intracortical ketamine in stressed rats (50). Conversely, higher doses (> 20 mg/kg) and/or long-term administration (6 months) of ketamine impairs working memory in non-stressed mice (98, 99), indicating that its cognitive effects may depend on factors such as drug dosage, treatment duration, and prior exposure to stress. Our findings also suggest that ketamine and BSM reverse CUS-induced deficits in a mouse paradigm of social approach. Loss of social interaction after chronic restraint stress or CUS was reversed by the α5-GABA_A_R NAMs MRK-016 or L-655,708 within 24 h in rats (33). However, in the rat social approach–avoidance test, BSM administration in non-stressed rats did not improve preference for the social compartment containing an unfamiliar conspecific (36), suggesting that stress history, including the chronic social isolation used here, may sensitize social circuits to the effects of these α5-NAMs.

At the circuit-level, strategies that target the GABAergic and glutamatergic systems provide more direct and fast control over neuronal activity and circuit modulation (14, 19, 39, 57, 79). In the mPFC, either direct photostimulation of pyramidal cells or indirect disinhibition of these neurons through chemogenetic silencing of GABA interneurons, including SST and PV subtypes, is sufficient to trigger ketamine-like rapid antidepressant actions (39, 63). Alongside this rapid increase in glutamatergic activity, our recent calcium imaging recordings in freely behaving mice provide evidence that ketamine exerts sustained antidepressant-like effects through a delayed and long-lasting facilitation of GABAergic function, which is engaged *during* behavioral testing 24–72 h after treatment (13). Therefore, we hypothesized that this temporal dynamics of glutamate and GABA activity may also underlie the behaviors facilitated by α5-NAMs. Consistent with this idea, chemogenetic silencing of mPFC CaMKIIα^+^ neurons completely blocked BSM-induced behavioral responses at all tested timepoints. Interestingly, while silencing of mPFC GABA interneurons did not reverse the short-term effects of BSM, it fully abolished its long-term behavioral responses. Taken together, these findings support the idea that BSM enhances excitatory activity and downstream intracellular signaling by reducing α5-GABA_A_R-mediated inhibitory tone on mPFC pyramidal neurons, with GABAergic adaptations contributing to the fine-tuning of long-term plasticity (Fig. 6F).

As a limitation of this study, the chemogenetic approach did not allow for temporally precise monitoring of neuronal activity underlying the behavioral effects of BSM, nor investigated the role of specific glutamatergic and GABAergic neuronal subpopulations (e.g., Drd1 vs. SST neurons). These questions will be addressed in future studies, for which approaches such as optogenetic manipulation and fiber photometry–based calcium recordings will be particularly valuable to resolve the temporal dynamics of E/I activity following treatment. Future work will also be important to determine how hippocampal connectivity and sex-dependent differences contribute to these effects. Notwithstanding these methodological considerations, our results contribute to a broader knowledge on the mechanism of action of BSM to produce antidepressant, procognitive and prosocial effects, with α5-NAMs representing a promising strategy for development of novel MDD pharmacotherapies.

## Acknowledgements

We thank Dr. Marina Picciotto for providing the Cre-expressing mouse strains; Samantha Colayori for the technical assistance in Fogaça Lab; members of Fogaça Lab for assisting the first author with the chronic unpredictable stress protocol; Brian I. Knapp for the technical assistance with experiments of conditioned place preference; the staff of the Vivarium and Division of Comparative Medicine (DCM), University of Rochester, for the specialized training and animal care. This study was supported by grant R00MH126098 (NIMH) and NARSAD Young Investigator Award (29063, BBRF Foundation).

## Author contributions

F.D. and M.V.F. designed the experiments. F.D. and M.V.F. performed the mouse studies, analyzed all datasets, and generated all figures. C.T.F. assisted M.V.F. with c-Fos experiments and assisted F.D. with viral microinjections in transgenic mice. J.M.B. assisted with the CPP test and reviewed the manuscript. O.P. and J.A. synthesized and provided BSM and reviewed the manuscript. F.D. wrote the manuscript, and M.V.F. reviewed and edited it. M.V.F. acquired funding, conceived the study, and supervised the project. All authors contributed comments on the final manuscript.

