## Supplemental Figures for "Negative allosteric modulation of α5-GABA_A_ receptors engages dynamic cortical glutamatergic and GABAergic mechanisms underlying adaptive behavior in mice"

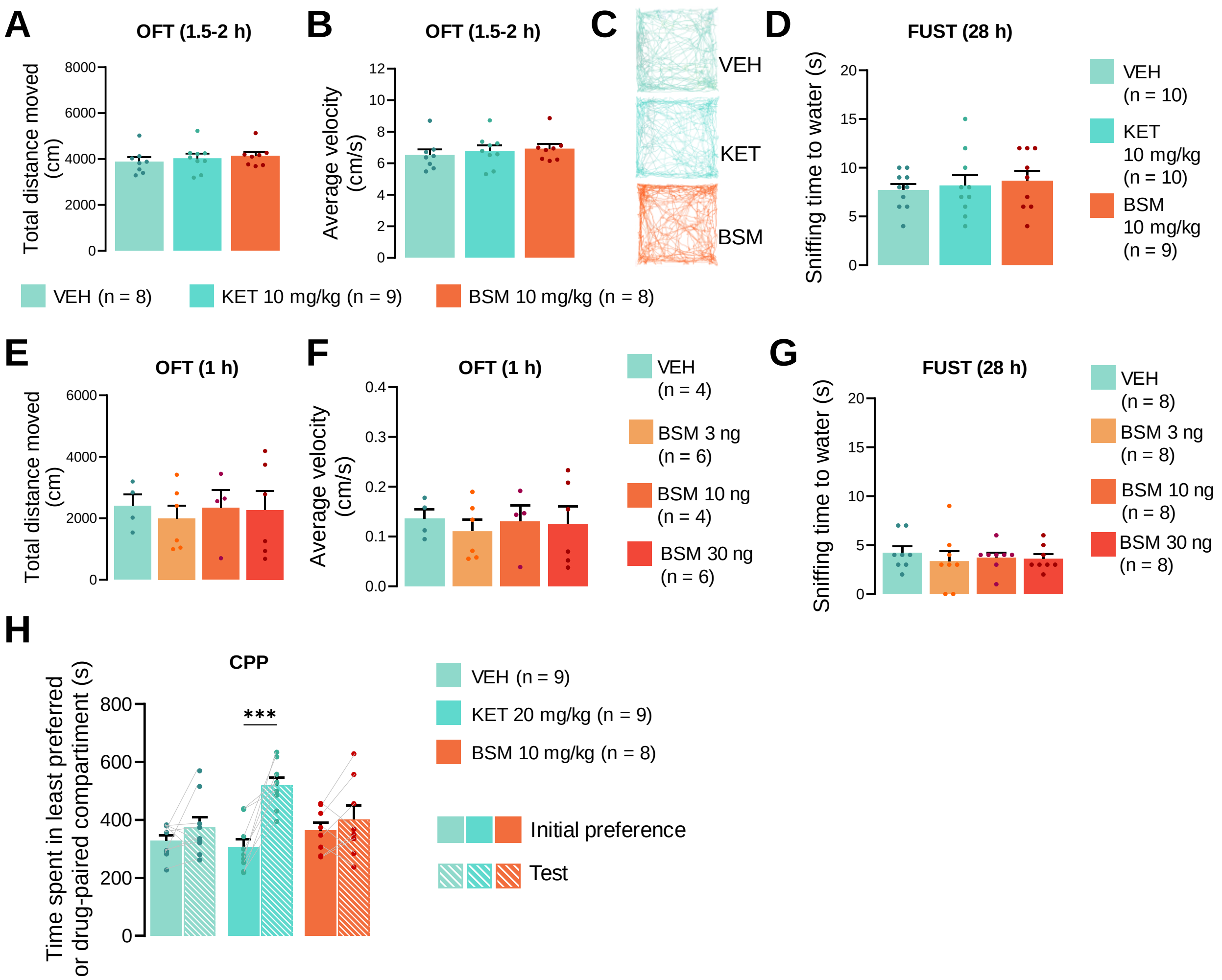


**Figure S1**. **Basmisanil (BSM) lacks locomotor and reinforcing effects.** (**A**) Average total distance (F(2,22) = 0.42; P=0.66) and (**B**) velocity (F(2, 22) = 0.35; P = 0.71) moved during the open field test (OFT) 1.5-2 h post-treatment with vehicle (VEH), ketamine (KET, 10 mg/kg i.p.) or BSM (10 mg/kg, i.p.). (**C**) Locomotion traces of mice exploring the open field arena for each group. (**D**) Average time spent sniffing a cotton tip dipped in water during the female urine sniffing test (FUST, F(2,26) = 0.28, P = 0.76). (**E**) Average total distance (F(3, 16) = 0.12; P = 0.95) and (**F**) velocity F(3, 16) = 0.14; P = 0.94) moved during the OFT 1 h following BSM microinjection (3, 10 or 30 ng) into the mPFC. (**G**) Average time spent sniffing a cotton tip dipped in water during the FUST following intra-mPFC microinjection (F(3, 29) = 0.56; P = 0.65). Data are plotted as mean ± SEM. P values were calculated using one-way ANOVA or two-sided paired Student’s t-test. ***P ≤ 0.001.

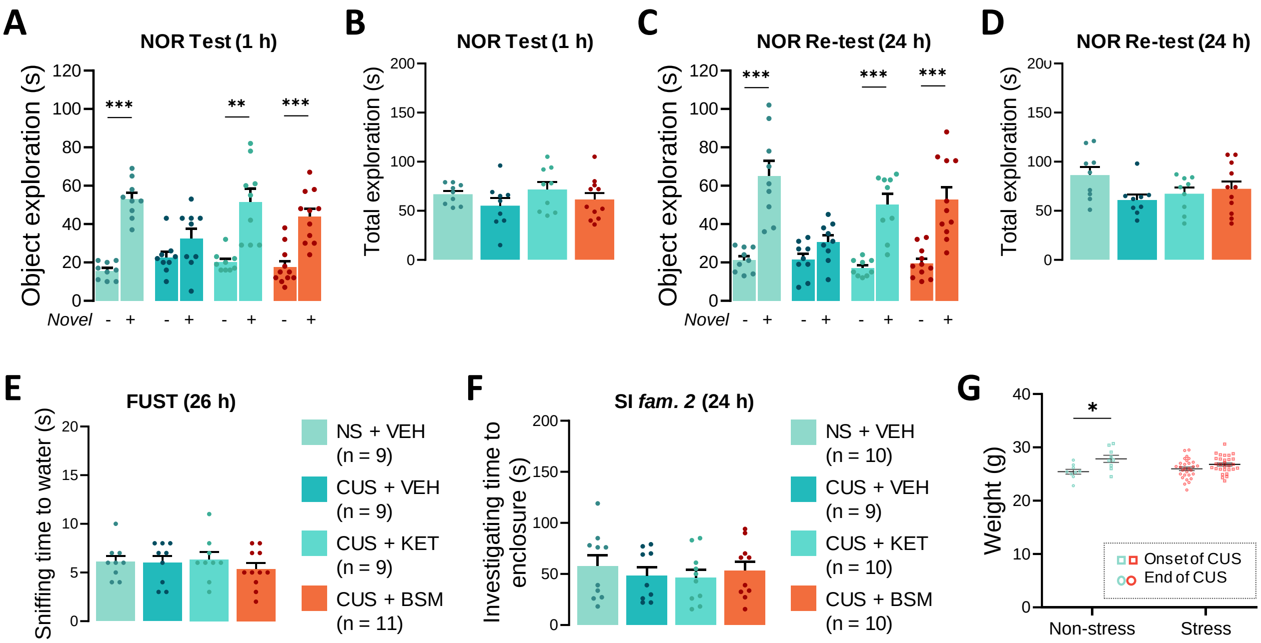


**Figure S2. Effect of Basmisanil (BSM) or ketamine (KET) on object exploration, water sniffing, empty-enclosure investigation, and body weight gain after the chronic unpredictable stress (CUS) protocol.** (**A**) Time spent exploring familiar and novel objects during the test session performed 1 h post-treatment with vehicle (VEH), ketamine (KET, 20 mg/kg, i.p.) or BSM (10 mg/kg, i.p.; Non-stressed, NS + VEH: t(16) = -10.10, P < 0.001. CUS + VEH: t(12.90) = -1.66, P = 0.12. CUS + KET: t(9) = -4.44, P = 0.002. CUS + BSM: t(18) = -4.90, P < 0.001). (**B**) Total time spent exploring both familiar and novel objects 1 h post-treatment (F(3,34) = 1.1, P = 0.35). (**C**) Time spent exploring familiar and novel objects during the re-test session 24 h post-treatment (NS + VEH: t(9.18) = -5.44; P < 0.001. CUS + VEH: t(16) = -1.97; P = 0.07. CUS + KET: t(9.17) = -5.94; P < 0.001. CUS + BSM: t(12.84) = -4.94; P < 0.001). (**D**) Total time spent exploring both familiar and novel objects 24 h post-treatment ($F(3, 34) = 2,16$, $P = 0.11$). (**E**) Average time spent sniffing a cotton tip dipped in water during the FUST ($F(3,34) = 0.42$, $P = 0.74$) and (**F**) investigating an empty enclosure during the SI test ($F(3, 35) = 0.33$, $P = 0.80$). (**G**) Body weight gain in NS ($t(8) = 2.7$; $P = 0.025$) and stressed ($t(24) = 1.7$; $P = 0.10$) male mice at the end of the experiment compared with the onset of CUS. Data are plotted as mean ± SEM. P values were calculated using one-way ANOVA or two-sided Student's t test. $^{*}P \leq 0.05$; $^{**}P \leq 0.01$; $^{***}P \leq 0.001$.

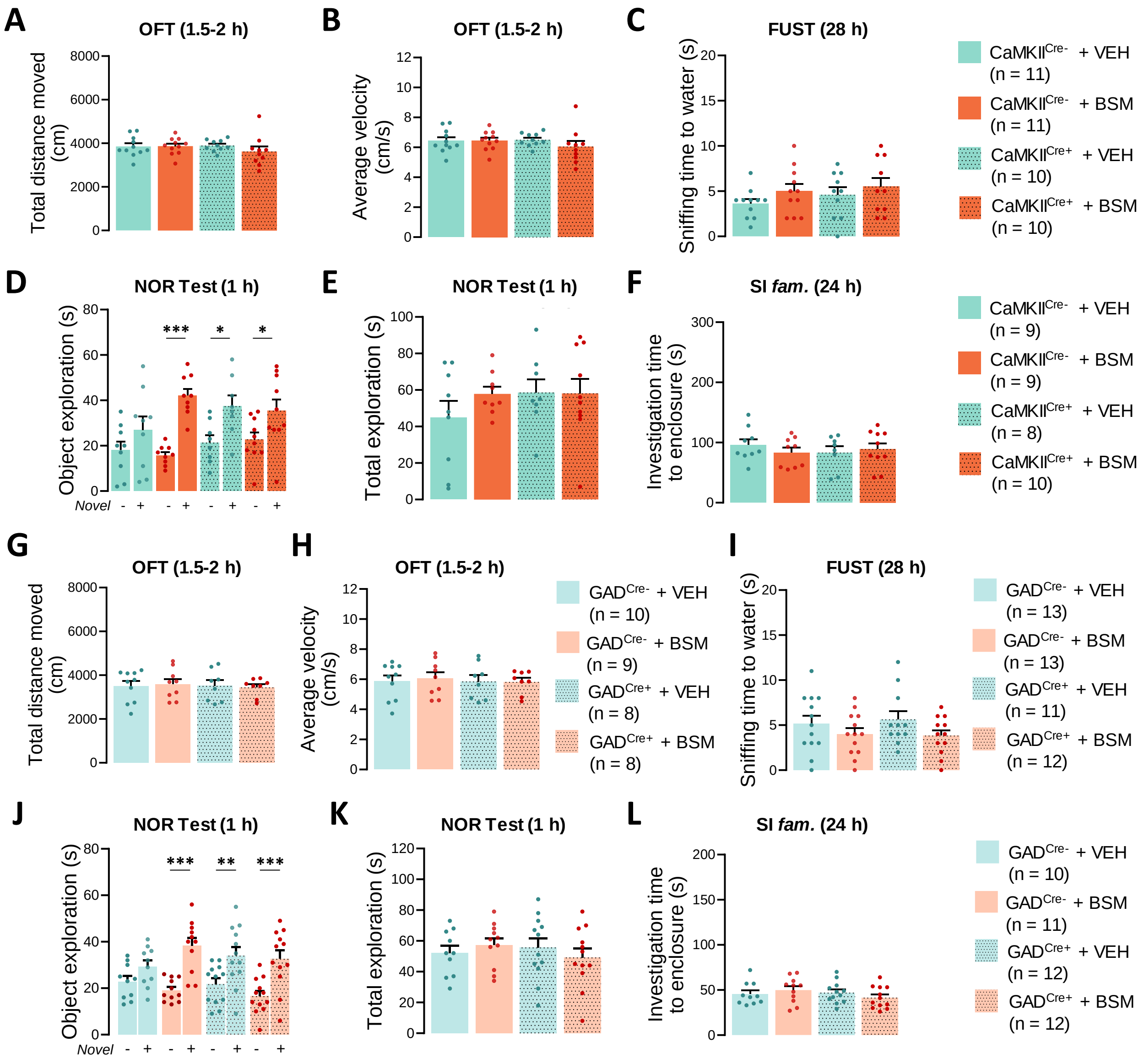

**Figure S3. Effect of Basmisanil (BSM) on novel object exploration, locomotor activity, water sniffing and empty-enclosure investigation in CaMKII$^{Cre+}$ and GAD$^{Cre+}$ mice.** (**A**) Average total distance ($F_{int}$ (1,38) = 0.91, P = 0.34; $F_{gen}$ (1,38) = 0.47, P = 0.50; $F_{treat}$ (1,38) = 0.75, P = 0.39) or (**B**) velocity ($F_{int}$ (1,38) = 0.96, P = 0.33; $F_{gen}$ (1,38) = 0.45, P = 0.51; $F_{treat}$ (1,38) = 0.78, P = 0.38) moved during the open field test (OFT) 1.5-2 h post-treatment with CNO (1 mg/kg, i.p.) and vehicle (VEH) or BSM (10 mg/kg, i.p.) in CaMKII$^{Cre}$ mice. (**C**) Average time spent sniffing a cotton tip dipped in water during the FUST ($F_{int}$ (1,38) = 0.09, P = 0.77; $F_{gen}$ (1,38) = 0.89, P = 0.35; $F_{treat}$ (1,38) = 2.13, P = 0.15). (**D**) Time spent exploring familiar *versus* novel objects during the test session 1 h post-treatment (CaMKII$^{Cre-}$ + VEH: t(14) = -1.17, P = 0.26. CaMKII$^{Cre-}$ + BSM: t(16) = -7.90, P < 0.001. CaMKII$^{Cre+}$ + VEH: t(10) = -2.84, P = 0.02. CaMKII$^{Cre+}$ + BSM: t(18) = -2.15, P = 0.05). (**E**) Total time spent exploring both familiar and novel objects ($F_{int}$ (1,31) = 0.77, P = 0.38; $F_{gen}$ (1,31) = 0.85, P = 0.36; $F_{treat}$ (1,31) = 0.82, P = 0.37). (**F**) Average time spent investigating an empty enclosure during the SI test ($F_{int}$ (1,31) = 0.46, P = 0.50; $F_{gen}$ (1,31) = 0.003, P = 0.96; $F_{treat}$ (1,31) = 0.004, P = 0.95). (**G**) Average total distance ($F_{int}$ (1,31) = 0.11, P = 0.74; $F_{gen}$ (1,31) = 0.10, P = 0.75; $F_{treat}$ (1,31) = 0.001, P = 0.98) or (**H**) velocity ($F_{int}$ (1,31) = 0.08, P = 0.77; $F_{gen}$ (1,31) = 0.11, P = 0.75; $F_{treat}$ (1,31) = 0.04, P = 0.84) moved during the OFT 1.5-2 h post-treatment in GAD$^{Cre}$ mice. (**I**) Average time spent sniffing a cotton tip dipped in water during the FUST ($F_{int}$ (1,45) = 0.17, P = 0.68; $F_{gen}$ (1,45) = 0.04, P = 0.84; $F_{treat}$ (1,45) = 3.59, P = 0.06). (**J**) Time spent exploring familiar *versus* novel objects during the test session 1 h post-treatment (GAD$^{Cre-}$ + VEH: t(18) = -1.80, P = 0.09. GAD$^{Cre-}$ + BSM: t(20) = -5.21, P < 0.001. GAD$^{Cre+}$ + VEH: t(22) = -2.74, P = 0.01. GAD$^{Cre+}$ + BSM: t(22) = -3.75, P = 0.001). (**K**) Total time spent exploring both familiar and novel objects ($F_{int}$ (1,41) = 1.22, P = 0.28; $F_{gen}$ (1,41) = 0.17, P = 0.68; $F_{treat}$ (1,41) = 0.01, P = 0.91). (**L**) Average time spent investigating an empty enclosure during the

SI test ($F_{int}$ (1,41) = 1.70, P = 0.20; $F_{gen}$ (1,41) = 0.83, P = 0.37; $F_{treat}$ (1,41) = 0.03, P = 0.87). Data are plotted as mean ± SEM. P values were calculated using two-way ANOVA or two-sided Student's t test. *P ≤ 0.05; **P ≤ 0.01; ***P ≤ 0.001.
